# Fast and Comprehensive N- and O-glycoproteomics analysis with MSFragger-Glyco

**DOI:** 10.1101/2020.05.18.102665

**Authors:** Daniel A. Polasky, Fengchao Yu, Guo Ci Teo, Alexey I. Nesvizhskii

## Abstract

Glycosylation is a ubiquitous and heterogeneous post-translational modification (PTM) used to accomplish a wide variety of critical cellular tasks. Recent advances in methods for enrichment and mass spectrometric analysis of intact glycopeptides have produced large-scale, high-quality glycoproteomics datasets, but interpreting this data remains challenging. In addition to being large, complex, and heterogeneous, glycans undergo fragmentation during vibrational activation, making common PTM search strategies ineffective for their identification. We present a computational tool called MSFragger-Glyco for fast and highly sensitive identification of N- and O-linked glycopeptides using open and glycan mass offset search strategies. Reanalysis of recently published N-glycoproteomics data resulted in annotation of 83% more glycopeptide-spectrum matches (glycoPSMs) than in previous results, which translated to substantial increases in the numbers of glycoproteins and glycosites that could be identified. In published O-glycoproteomics data, our method more than doubled the number of glycoPSMs annotated when searching the same peptides as the original search and resulted in up to a 6-fold increase when expanding searches to include large numbers of possible glycan compositions and other modifications. Expanded searches revealed trends in glycan composition and crosstalk with phosphorylation that remained hidden to the original search. With greatly improved spectral annotation, coupled with the fast speed of fragment ion index-based scoring, MSFragger-Glyco makes it possible to comprehensively interrogate glycoproteomics data and illuminate the many roles of glycosylation.

## Introduction

Glycosylation is a ubiquitous and heterogeneous post-translational modification (PTM) of proteins used by cells to accomplish a wide variety of critical tasks and provide a flexible response to a changing environment^1,2^. Altered glycosylation profiles have been detected or implicated numerous cancers and other diseases, making the comprehensive characterization of protein glycosylation critical to understanding health and disease^3–5^. Analysis of intact glycopeptides by tandem mass spectrometry (MS) has the potential to simultaneously determine the sites and compositions of glycans on a proteome-wide scale, but presents several challenges due to the unique characteristics of glycosylation. Enrichment of glycopeptides is required to overcome low ionization efficiencies in positive ion mode and the resulting ion suppression by peptides lacking glycosylation^6^. The heterogeneity of glycan compositions at a given site (micro-heterogeneity) and of the occupancy of possible glycosylation sites in a protein (macro-heterogeneity) present enormous challenges to interpretation of intact glycopeptide MS data^7,8^. Recent advances in enrichment and mass spectrometric analysis of intact glycopeptides have begun to produce large-scale, high-quality datasets from a range of organisms and sample types^9–13^. The ability to produce glycoproteomic data at this scale has the potential to generate a paradigm shift in understanding the role of glycosylation in health and disease.

Interpretation of proteome-scale intact glycopeptide mass spectrometry data remains challenging, however. The most common method for interpreting glycopeptide mass spectra is similar to the treatment of other PTMs in proteomics database searches, *i.e*., to search all or a subset of potential glycans as variable modifications on all possible glycosylation sites. Several existing proteomics search engines have been adapted to search glycosylation as a variable modification^14–16^, and several glycopeptide-specific tools tailored to particular MS acquisition methods also use variable glycan modifications for small-scale searches^17–20^. The variable modification approach has two major limitations: first, the large number of possible glycan masses can result in a combinatorial explosion of possibilities for peptides that contain multiple potential glycosylation sites, which is particularly problematic for the analysis of peptides with densely clustered O-linked glycans. The second is the highly labile nature of glycosylation during vibrational activation, including the collisional activation used solo or in a hybrid mode in the vast majority of glycopeptide analyses. The variable modification approach as employed by many tools, *e.g*. Byonic^15^ and SEQUEST^16^, looks for fragment ions containing the intact glycan, even though glycan fragmentation during collisional or hybrid activation makes these ions unlikely to appear. Some search engines, *e.g*. Comet^21^, allow neutral losses to be specified for labile modifications, but the diversity of possible glycans would result in a prohibitive number of neutral loss masses.

An alternative approach can be found in the open search method^22–29^. In open searches, the peptide mass is determined by matching fragment ions without knowledge of the precursor, and the difference between the matched sequence mass and the precursor mass, called the “delta mass” or “mass offset,” is the mass of any unspecified modifications to the sequence. Crucially, for modifications that are labile, this strategy captures their presence on the precursor via the delta mass without requiring their presence on fragment ions, enabling a larger proportion of the observed fragment ions to be matched. A subset of open search, called a mass offset^30^ or multinotch^23^ search, uses this strategy to look only for a known set of delta masses of interest, such as those of known or potential glycans. A recently introduced glycoproteomics search tool, pGlyco 2.0^8^, takes advantage of this strategy in their “coarse scoring” approach, searching for delta masses that match entries in a glycan database and scoring those spectra on the presence of Y-type ions before sending candidates to peptide sequence searching. This two-step approach, however, requires that glycans generate abundant Y ion fragments, limiting its applicability to N-glycans fragmented by CID/HCD.

Here we present MSFragger-Glyco, a search engine that applies the concept of open and mass offset search strategies, made practical for proteomics applications using the fragment ion indexing approach of MSFragger^22^, to searching glycoproteomics data. Importantly, spectra are searched for all fragment ion series of interest simultaneously, including any of Y, *b/y* with no glycan or (optionally) with a single HexNAc remaining, and *c/z* ions containing the intact glycan, depending on the activation method(s) employed. This ensures that the score associated with any spectrum is generated from all fragment ions that can be reasonably expected, without noise from highly unlikely fragments, resulting in greatly improved confidence for the identification of labile glycans. Taking advantage of the ultrafast indexed-based searching, complex searches including hundreds of possible glycan masses can now be accomplished in a matter of minutes to hours. We applied MSFragger-Glyco to several state-of-the-art glycoproteomics datasets of N- and O-linked glycopeptides, comparing against published glycoPSM identifications from Byonic and SEQUEST. In all cases, the glycan mass offset strategy of MSFragger-Glyco provided a substantial increase in the number of spectra that could be successfully annotated, and corresponding increases in the numbers of glycopeptides, proteins, and sites identified that could be identified from the data. For O-glycoproteomics in particular, this approach offered 2-6-fold improvements in glycoPSMs identified over recently published results from the same raw data, indicating the potential of MSFragger-Glyco for widespread improvement in the analysis of glycoproteomic data.

## Results

### Development of MSFragger-Glyco

MSFragger-Glyco takes advantage of the localization-aware (i.e. including shifted fragment ions) open search strategy^31^, with several modifications specific to glycopeptides (Fig. 1). Fragmentation of glycopeptides results in a complex milieu of products, especially in hybrid activation techniques such as EThcD and AI-ETD^10,11,32–35^. Because fragmentation of glycans is typically lower energy than that of the peptide backbone during CID/HCD, it is unusual to observe the intact mass of the glycan on any fragment ions, except in ETD/ECD without supplemental activation. This presents a challenge to typical (variable modification) searches, in that the observed mass of the precursor no longer matches that of the sequence that can be detected from the fragment ions. The resulting difference between the peptide sequence and the precursor mass is referred to as “mass offset”. MSFragger-Glyco can perform a fully open search (i.e. allow any mass offset) for exploratory interrogation of the data, and improved performance can be achieved by restricting allowed mass offsets a set of user-provided glycan masses. An arbitrary number of these glycan masses can be supplied to MSFragger-Glyco and set to correspond to either N- or O-glycans. To improve glycopeptide searches, MSFragger-Glyco considers additional ion types, including Y ions (intact peptide plus any of several common glycan fragments) and *b/y* ions with a single HexNAc residue remaining. In addition, it performs sequence motif checks for peptides and an oxonium ion check for spectra. For each peptide that contains a potential glycosite (N-X-S/T for N-glycans or S/T for O-glycans by default), Y and (optionally) *b/y* + HexNAc ions are added to the fragment index (Fig 1a, right). The glycan mass offsets are only searched for peptides that contain a potential glycosite and for spectra that contain sufficiently abundant oxonium ions, *i.e*. those with evidence of glycan fragmentation. For all other peptides and spectra, a regular search is performed with no mass offsets or glycan masses allowed (Fig. 1b, left). Depending on the activation method employed, the glycan mass offset search can look for CID/HCD-type fragmentation: *b,y,Y*, and *b/y* + HexNAc ions, ETD/ECD-type fragmentation: *c/z* ions with the intact glycan, or hybrid fragmentation including all aforementioned ion types (Fig. 1B, right).

**Figure 1.**
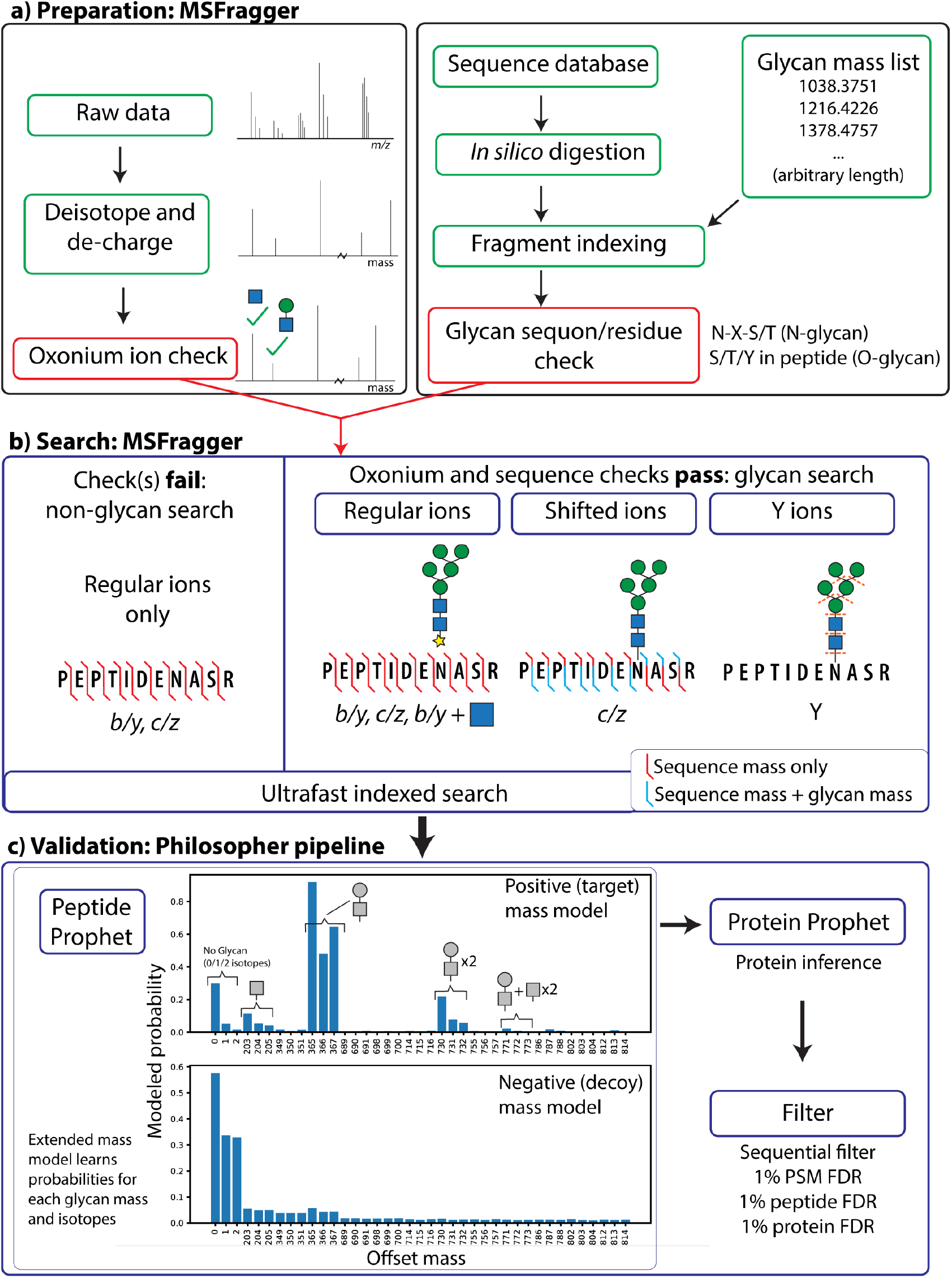
Workflow of MSFragger-Glyco. a) Data and database preparation. Raw MS/MS spectra are deisotoped, de-charged, and checked for the presence of oxonium ions. Protein sequences are digested *in silico* and fragment ions are indexed. Peptides containing a possible glycosite have additional glyco-specific fragments added to the index. b) Spectra are searched against indexed peptides. If the spectrum contains oxonium ion(s) and the peptide being considered contains a possible glycosite, a glycan search is performed (right); if either check fails, a regular search is performed. Shifted ions (blue) contain the intact mass of the glycan on the peptide while regular ions (red) contain only the masses of the amino acid residues. c) FDR filtering is performed using Philosopher. Plot of probabilities learned by PeptideProphet for mass offsets from O-glycopeptide data shows high probability for common compositions (e.g. HexNAcHex at 365) and isotopes in the positive model, whereas the negative model shows similar probability for across mass shifts.

Potential peptide-spectrum matches (PSMs) are filtered using PeptideProphet^36^ and ProteinProphet^37^ using the Philosopher^38^ toolkit (https://philosopher.nesvilab.org/). We utilize the extended mass model of PeptideProphet to independently model peptides with different mass offsets^22^, corresponding to different glycan masses in this case. As a result, PSMs with similar database search scores may have very different modeled probabilities if, for example, one has a mass offset corresponding to a commonly observed glycan and the other the mass offset of a rare glycan. This effect can be seen in the output probability distributions for a subset of O-glycan masses searched (Fig. 1c, top), for example, in which a delta mass of 365 Da, corresponding to the very common HexNAc-Hex glycan, has very high probability while a delta mass of 349 Da, corresponding to the much less common HexNAc-Fuc glycan, has nearly zero probability in this data. Delta mass values of 366 and 367 Da also obtain high probabilities, as they occur when a non-monoisotopic precursor peak was selected, resulting in a delta mass of the glycan (365) plus precursor isotope error (1 or 2). The +2 isotope error (offset mass of 367, 732, etc.) appears to have elevated probability relative to the +1 isotope error because delta mass values are binned at a width of 1 Da in PeptideProphet, which can include multiple sub-populations, such as the combination of isotope +1 error and deamidation. The distribution of delta mass probabilities for decoys (Fig. 1c, “negative”) shows roughly even probabilities for all glycan delta mass values, as hits to decoy peptides are expected to occur randomly without enrichment for specific glycan compositions. Following PeptideProphet, protein inference is performed using standard open search settings in ProteinProphet and filtering is performed in Philosopher to 1% PSM and protein levels. A sequential filtering step is then applied to remove any PSMs matched to proteins that did not pass 1% protein-level FDR.

### MSFragger-Glyco Greatly Improves Identification of Labile Glycan Spectra

To demonstrate the utility of MSFragger-Glyco for large-scale glycoproteomics analyses, we searched several publicly available datasets. Riley *et al*^10^ recently published a large-scale analysis of N-glycosylation of mouse brain tissue using hybrid activation method AI-ETD, representing the largest number of glycosylation sites found in such tissue to date, including many sites not previously observed or confirmed by experimental evidence. To analyze the data, Riley *et al*. used Byonic^15^, a commercial platform that uses a variable modification-type search with tunable control of “common” and “rare” modifications, to handle glycosylation analyses. As with other variable modification-type searches, Byonic places potential N-glycans on peptides containing possible glycosylation sites and looks for fragments of those peptides that contain the intact glycan.

A detailed comparison reveals substantial benefits of the MSFragger-Glyco mass offset search over this more typical variable modification strategy. An example MS/MS spectrum of a glycopeptide selected for HCD fragmentation, shown in Fig. 2a, is dominated by Y and B (oxonium) ions resulting from fragmentation of the glycan, while only a small fraction of the ion current comes from fragmentation of the peptide backbone, typically following extensive or complete fragmentation of the glycan. Peaks that would be considered in a variable modification search, *i.e*. peaks explained by the peptide sequence with an intact glycan present on Asn-9, are shown in red, and peaks matched by MSFragger-Glyco are shown in blue. By matching glycans as mass offsets between observed sequence and precursor masses instead of direct modifications, any peptide fragments that contain the original position of the glycan can be matched successfully (light blue), whereas none are matched in the variable modification search because none are observed with the intact glycan. In addition, MSFragger-Glyco adds the Y (dark blue) and *b/y* + HexNAc (medium blue) ion series to the mass offset search for glycopeptides, which represent the majority of matched ions in this spectrum. As a result, the conventional variable modification search matches 8 ions and <5% of the total ion current, while with the MSFragger-Glyco’s mass offset search with glycan-specific ion types matches 21 ions and >50% of the total ion current, generating a far more confident PSM. For AI-ETD spectra, glycan fragmentation from the laser irradiation results in a similar effect, although typically less so than for HCD since the degree of glycan fragmentation is lower, and some *c/z*-type ions can be observed with the glycan intact^10^. Importantly, the MSFragger-Glyco glycan offset search also includes shifted ions, which contain the intact glycan for *c/z*-type fragments, allowing for matching of all ion types observed in AI-ETD and other hybrid methods, such as EThcD.

**Figure 2.**
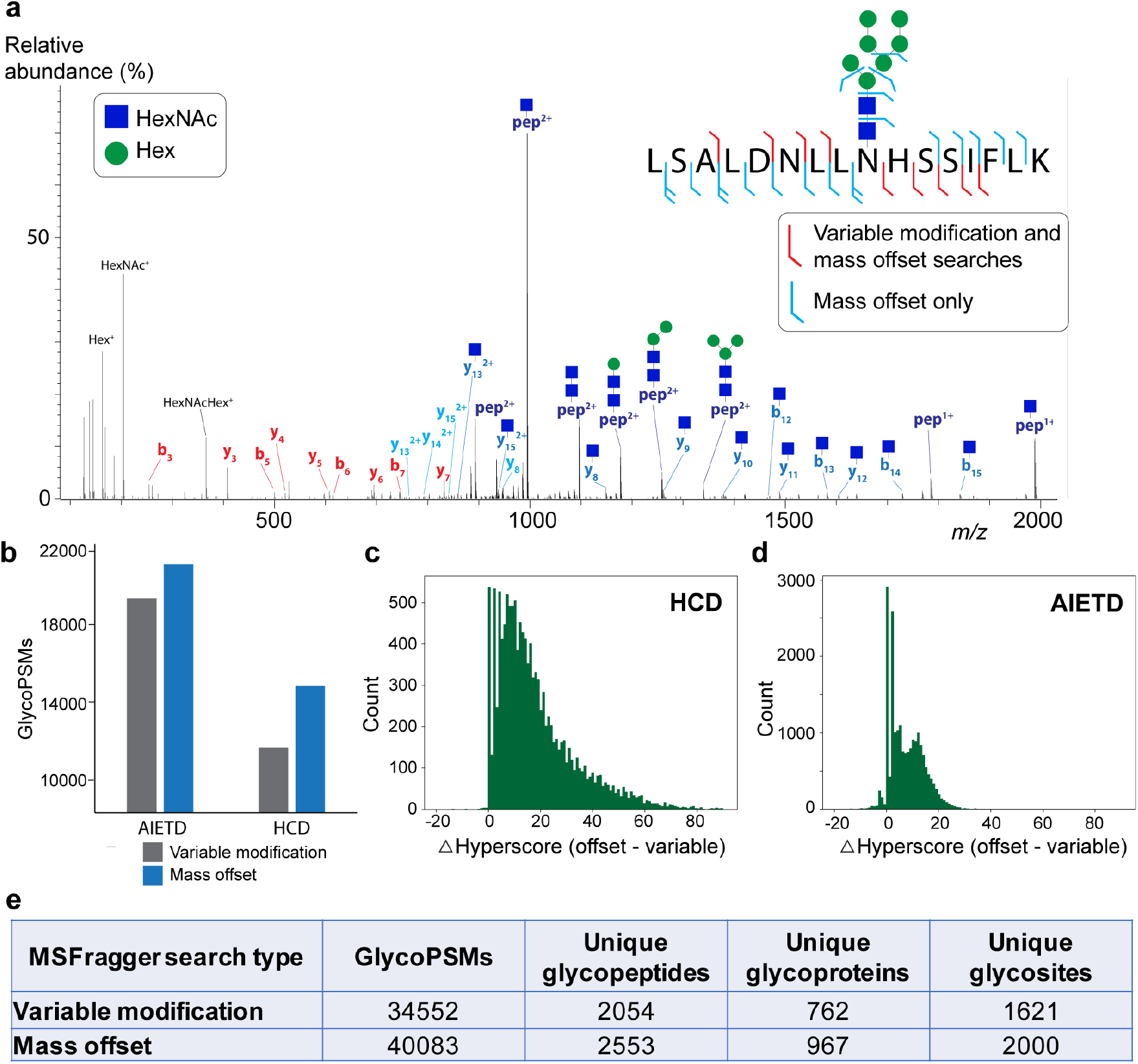
Comparison of mass offset and variable modification-type searches for N-linked glycopeptides. a) Example HCD tandem mass spectrum of peptide LSALDNLLNHSSIFLK with glycan HexNAc2Hex7. Fragment ions that match the identification assuming the intact glycan is present at N-9 (variable modification-type) are colored red. Note that none of the fragments annotated in red contain the glycosite. Fragments in blue correspond to the mass offset search, including Y ions, *b/y* ions without the glycan or with a single HexNAc (blue square) remaining. Oxonium ions are shown in black (not all are labeled). b) Number of glycoPSMs obtained for MSFragger mass offset search (orange) or variable modification search (blue) from AIETD and HCD spectra. c) Score difference (mass offset hyperscore – variable modification hyperscore) for spectra that were annotated in both search types for HCD spectra and (d) for AI-ETD spectra, showing a much larger improvement in scores for HCD data. e) Table of results for MSFragger-Glyco mass offset and variable modification searches of 17 glycans.

To compare the two search types across the complete dataset, we performed a variable modification search with MSFragger, in which 17 commonly occurring glycans were specified as variable modifications on the consensus sequon N-X-S/T. An MSFragger-Glyco search with the same 17 glycans specified as mass offsets was performed with the same protein database and all other parameters, leaving the different fragment ions considered as the only difference between searches. For both HCD and AI-ETD activation, the mass offset search annotated many more glycoPSMs than the variable modification search and, as expected, the degree of improvement was larger in HCD spectra (24% increase) than AI-ETD (8% increase) (Fig. 2b). In many cases, the both searches successfully identified a glycoPSM, but with very different levels of confidence. For HCD spectra, nearly all glycoPSMs (>95%) had a higher score in the mass offset search, with over 60% having a substantial increase of more than 10 (Fig. 2c). As expected, the effect was less pronounced in AI-ETD spectra, but 81% of all glycoPSMs still scored higher in the mass offset search, with 33% having an increase of more than 10 (Fig. 2d). Overall, the increased scores and confidence of the mass offset search resulted in nearly 5,000 more glycoPSMs than in the variable modification search, translating to 20-25% increases in the number of unique glycopeptides, glycoproteins, and glycosites observed in the data (Fig. 2e). The ability of the mass offset search to capture the expected fragment ions from glycopeptides thus gives it a unique advantage over traditional variable modification searches, resulting in increases in number of spectra identified and their confidence, and ultimately of identified glycopeptides and glycoproteins.

### Large-scale N-glycoproteomics with MSFragger-Glyco

To evaluate the performance of the MSFragger-Glyco mass offset method for N-glycoproteomic data, we searched the AI-ETD data from Riley *et al*. using the same 182 possible glycan compositions and protein database used in their search, as well as the same digestion and non-glycan modification parameters. MSFragger-Glyco obtained a dramatic increase in the number of spectra that can be successfully matched to glycopeptides, with 44,187 glycoPSMs to the 24,099 reported in Riley *et al* (Fig. 3a). This increase in identified spectra translated to a 57% increase in the total number of unique glycopeptide sequences detected, and a 38% increase in unique glycoproteins and glycosylation sites identified across the entire dataset (Fig. 3a). The complete MSFragger-Glyco search, including spectral pre-processing and mass calibration, took 1.9 minutes per file on a desktop computer (6 cores, 32 GB RAM), which, to our knowledge, is substantially faster than existing tools. Compared to Riley *et al*. and Liu *et al*.^7^, another study with N-glycosylation data from mouse brain tissue, our analysis of the Riley *et al*. data shows excellent overlap with the previously detected glycosites while adding nearly 800 new glycosites not detected in either previous analysis (Fig. 3b). Both our MSFragger-Glyco search and Riley *et al*. report glycosites with a mixture of UniProt annotation levels, with the vast majority (>85% in both cases) lacking previous experimental evidence (Fig. 3c). In both cases, the majority of glycosites detected are predicted by sequence or similarity analyses in UniProt but without experimental evidence (Fig. 3c, light blue), with MSFragger-Glyco confirming 257 more predicted glycosites than the previous analysis. MSFragger-Glyco also detects 323 more potential glycosites with no UniProt annotation (Fig. 3c, orange). Despite the large increases in detected glycosites, the distribution of glycosites observed per glycoprotein and in glycan compositions observed per site remain very similar to those reported in Riley *et al*. (Fig. 3d, e).

**Figure 3.**
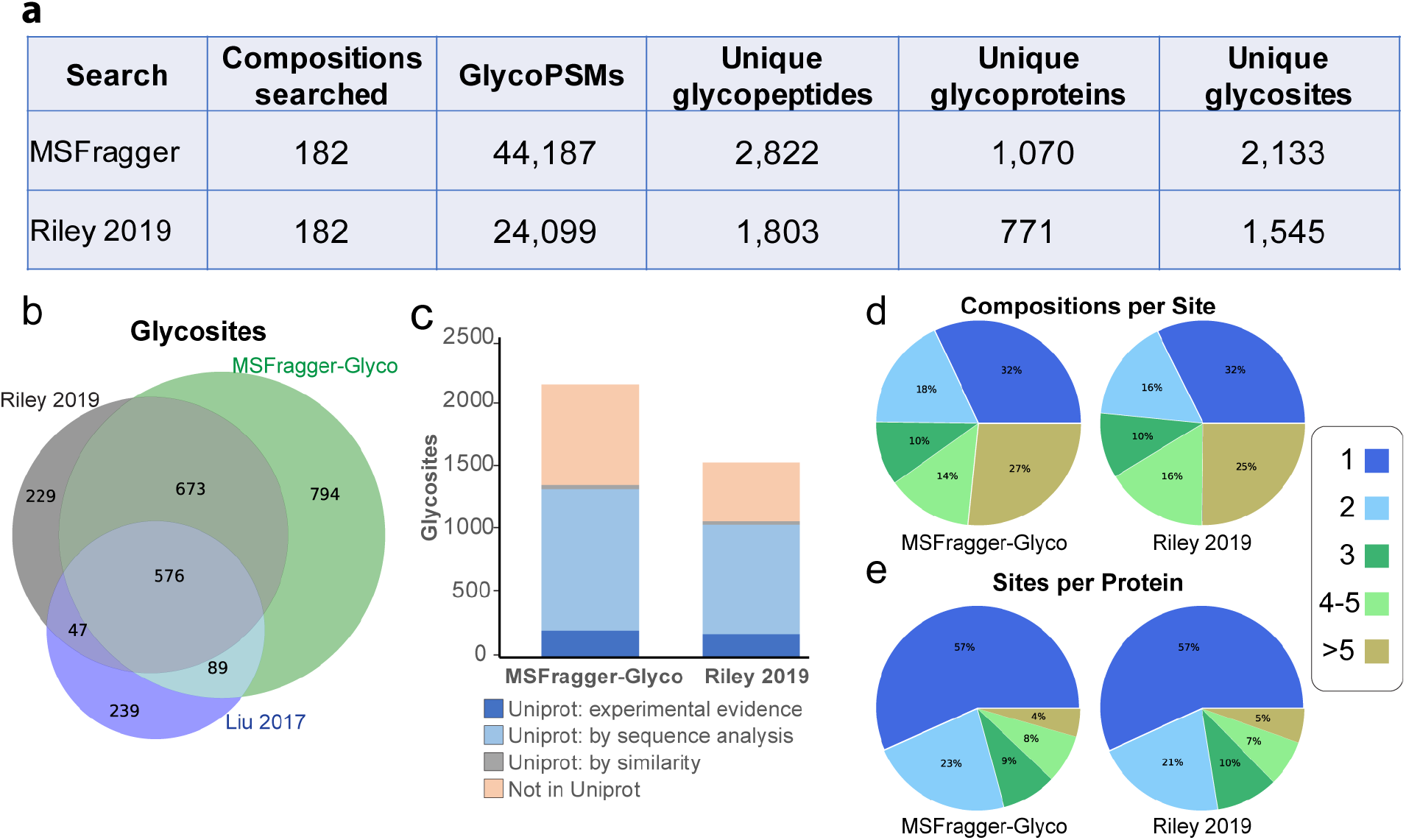
Comparison of MSFragger-Glyco and Riley *et al*. results. a) Direct comparison with identical protein databases and possible glycan compositions between MSFragger-Glyco and previously published Byonic results. b) Glycosite comparison between MSFragger-Glyco, Riley *et al*. and Liu *et al*. Note that MSFragger-Glyco and Riley *et al*. are generated from the same raw data; Liu *et al*. is a different dataset in the same tissue type. c) Comparison of found glycosites to UniProt, color coded by evidence type. d) Number of glycan compositions per glycosites and e) Number of glycosites per protein found in MSFragger-Glyco and Riley *et al*., showing very high similarity in both cases.

Overall, these results indicate that MSFragger-Glyco mass offset search performs exceptionally well for analyzing large-scale N-glycoproteomics data in HCD and hybrid activation modes. The 83% increase in glycoPSMs annotated from the same raw data with identical glycan and protein databases resulted in notable increases in glycoprotein and glycosite annotation, including confirmation of predicted glycosites and annotation of novel ones. With any large increase in reported PSMs, it is important to confirm that FDR control and validation was used appropriately to rule out inflation by low-confidence identifications, particularly given that FDR control in glycoproteomics remains challenging^39,40^. There are a number of key differences in our approach to FDR compared to Byonic, including use of data combined from all fractions and replicates for confidence modeling and protein inference. This method provides improved statistics, particularly for less common glycans that may only appear in a few PSMs in each individual dataset. Using the extended mass model in PeptideProphet allows us to model probabilities for each glycan mass individually, preventing aggregation of high and low-confidence modifications into a single FDR rate. To confirm that this was working as intended, we calculated the FDR for each glycan mass individually. There was not a significant difference between the FDR rates for glycoPSMs and non-glycoPSMs, and no obvious deviations from expected rates of decoy hits for any glycan masses (Supplementary Table 7). We also performed a search with an equal number of target and decoy glycans, with decoys generated by shifting the target masses by +20 Da, resulting in 0.6% of all PSMs matching to decoy glycans (Supplementary Table 3). Given these confirmations of our FDR approach and the vastly improved confidence in glycoPSMs observed in Fig. 2, the increases in our glycoPSM annotation rate can be attributed to the capability of the MSFragger-Glyco with mass offset method to capture peptide fragment ions following partial or complete glycan loss.

### Deciphering the vast complexity of O-glycoproteomics with MSFragger-Glyco

The several types of O-linked glycosylation are known to play important biological roles but have not been studied as extensively as N-linked glycosylation, due in part to additional challenges in enriching and analyzing O-linked glycans and glycopeptides. As in the case of N-glycosylation, O-linked glycans are highly labile during vibrational activation and occur in a wide variety of compositions. Unlike N-linked glycans, however, there is no consensus sequon for O-glycosylation, and some types of O-glycans are densely clustered in regions of protein sequence^41,42^. These factors make analysis of O-glycoproteomic MS/MS data particularly challenging, as many compositions potentially occurring on multiple sites within the same peptide results in a massive search space for any comprehensive O-glycan search by traditional methods. To date, large-scale O-glycoproteomics has largely proven intractable beyond simplified cases in which only certain types of O-glycans are generated, *e.g*. with SimpleCells^43^, by enzymatic reduction of glycan complexity^13^, or by searching for a small subset of highly abundant glycans in data that potentially contains many more compositions. These strategies, while effective in gleaning information from a very challenging problem, are unable to decipher the full complexity of O-glycosylation.

To evaluate the use of MSFragger-Glyco for large-scale O-glycoproteomics, we analyzed data from a recent study by Yang *et al.^12^*, which used a bacterial enzyme dubbed “OpeRATOR” that cleaves N-terminal to O-glycosylated Ser and Thr residues. The authors developed a protocol using this enzyme to enrich and analyze glycopeptides from kidney tissue, serum, and T-cells with HCD MS/MS. The data was searched using SEQUEST in a variable modification mode with two possible glycan types (masses) on Ser/Thr residues, resulting in the identification of nearly 35,000 glycoPSMs in the kidney tissue data (Table 1). This resulted in 12 total glycan compositions observed at the peptide level due to co-occurrence of glycans at multiple residues within some peptides. This is an important distinction when comparing with the mass offset search of MSFragger-Glyco, as the mass offset is computed at the peptide level, including all glycan modifications present on separate residues as a single mass. To compare search strategies, we performed both variable modification and mass offset searches in MSFragger-Glyco with the same two glycan types and search parameters. The variable modification search gave results similar to the SEQUEST search, finding 38,632 glycoPSMs. To perform the equivalent comparison with the mass offset method, we searched the 12 peptide-level compositions found in Yang *et al*. as mass offsets with MSFragger-Glyco, obtaining over 77,000 glycoPSMs, or more than double those found in the original search (Table 1). Even searching just two mass offsets (*i.e*. disallowing any peptides with multiple glycosites in the mass offset search) still resulted in a large increase, with over 50,000 glycoPSMs annotated. These increases are due in part to the high fragmentation energy used in the MS/MS analysis, resulting in very few fragment ions that retain a partial or intact glycan. The mass offset search is able to match these unmodified ions and use the offset between sequence and precursor masses to determine the glycan mass.

**Table 1.**
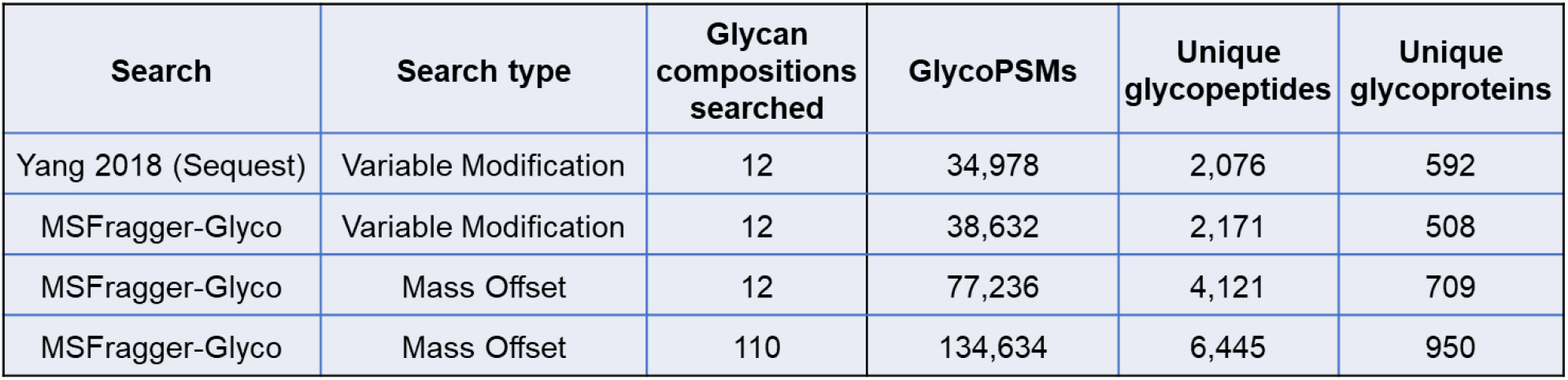
Comparison of MSFragger-Glyco and Yang *et al*. O-glycoproteomics search results in variable modification and mass offset modes. Note that variable modification searches with 2 glycan masses can identify multiple glycosites per peptide, resulting a total of 12 compositions found in Yang *et al*. The mass offset search used these same 12 glycan masses for comparison.

While the mass offset strategy considering equivalent modifications resulted in vastly more glycoPSMs than the previous searches, we sought to use the speed of MSFragger-Glyco to comprehensively analyze all glycopeptides present in the data. To do so, we first performed an exploratory, fully open search with MSFragger-Glyco to generate a list of abundant glycan compositions, then performed a mass offset search using this list. The open search revealed a large number of glycan compositions present in the data that were missed in the original searches, including fucosylated and sialylated glycans as well as masses corresponding to multiple glycans present on the same peptide. After removing compositions with overlapping isotope distributions, 110 total glycan compositions identified using open search were searched with the mass offset method, resulting in annotation of 134,634 glycoPSMs (Table 1), or 385% more than in the original search and nearly double the number from the 12-composition mass offset search. We found many additional glycopeptides, resulting in the identification of 358 more glycoproteins and many more potential glycosites than the original search of Yang *et al*. Several factors contributed to this massive increase in annotated spectra, including our expansion of the peptide search space as well as searching for many more types of glycans. Because OpeRATOR cleaves at glycosylated Ser/Thr, but the sites of glycosylation are not known in advance, Yang *et al*. digested their protein database by cleaving at all Ser/Thr residues but allowing up to 5 missed cleavages per peptide to allow for residues that may not be glycosylated. Taking advantage of MSFragger’s ultrafast indexed searching, we were able to allow up to 10 missed cleavages by OpeRATOR, and variable modifications including oxidation (M), guanidinylation and carbamidomethylation (K), phosphorylation (S, T, Y), deamidation (Q), and peptide N-terminal pyroglutamate formation (Q), after the exploratory open search revealed substantial amounts of these modifications in the data. These searches with very large peptide digestion and variable modification spaces, plus 110 potential glycan compositions, were still completed in a matter of hours, despite complexity that would be prohibitive for many search tools.

Given the proposed specificity of the OpeRATOR enzyme, Yang *et al*. assumed that all glycopeptides would contain a single glycosylation site at their N-terminus but noted the possibility that additional glycosites could be present if enzymatic cleavage at glycosylated Ser/Thr was imperfect. Our results show abundant evidence of these missed cleavages, particularly in cases where several glycosylated residues occur in series, for example in the peptide TTTIAEPDPGM from Glycophorin-C, which contains three known glycosylation sites in a row at the N-terminus (Supplementary Figure 1). The mass offset search annotates multiply glycosylated peptides as containing a single composite mass offset, which works well when the glycans have largely dissociated from peptide fragment ions. This approach is not ideal for determining the exact location of each glycosite, but given the relatively high energy at which the HCD fragmentation was conducted in this study, the majority of glycopeptide spectra lack fragment ions retaining any intact glycan or glycan fragments, precluding data-driven localization.

Applying the 110-composition search to the serum and T-cell datasets presented in Yang *et al*. yielded several interesting observations. In each case, the number of glycoPSMs annotated by our glycan mass offset search was dramatically increased compared to those reported by Yang *et al*., with 3.3 times as many glycoPSMs for the T-cell data and nearly 6.2 times as many glycoPSMs for the serum data (Table 2). The larger increase in PSMs in the serum samples can be attributed to the much greater proportion of fucosylated and sialylated glycans detected in serum (~33% of all PSMs were fucosylated and/or sialylated) as compared to kidney or T-cell samples (<15% of all PSMs fucosylated and/or sialylated), as the original search by Yang *et al*. did not consider fucosylated or sialylated glycans (Fig. 4). We also observed substantial co-occurrence of phosphorylation with O-glycosylation, particularly in the serum data, in which 18% of all glycoPSMs also contained phosphorylation (Fig. 4C). Crosstalk between phosphorylation and O-glycosylation is thought to play important roles in several cellular processes, but challenges searching both PTMs in proteomics data have largely prevented deeper understanding of these processes^44,45^. Indeed, the glycans detected on phosphorylated peptides contained a much higher proportion of complex fucosylated and sialylated glycans (nearly 60% of all glyco+phospho PSMs) than non-phosphorylated glycopeptides (Fig. 4C, right). While this complexity presents a significant challenge to traditional search tools, these results indicate that MSFragger-Glyco is well-suited to further investigation of this complex crosstalk. Yang *et al*. also highlight differences in glycosylation sites and occupancy between tumor and normal kidney tissue samples, focusing on Versican and Aggrecan core proteins. We find many additional glycoPSMs that broadly support the conclusion that glycosylation is increased in the tumor data, but exercise caution in localizing the glycans given the general lack of glycan-containing fragment ions in this dataset. There were no significant differences in observed glycan compositions between the normal and tumor kidney tissues (Fig. 4b) despite the clear increases in glycoPSMs and sites from the tumor samples.

**Table 2.**
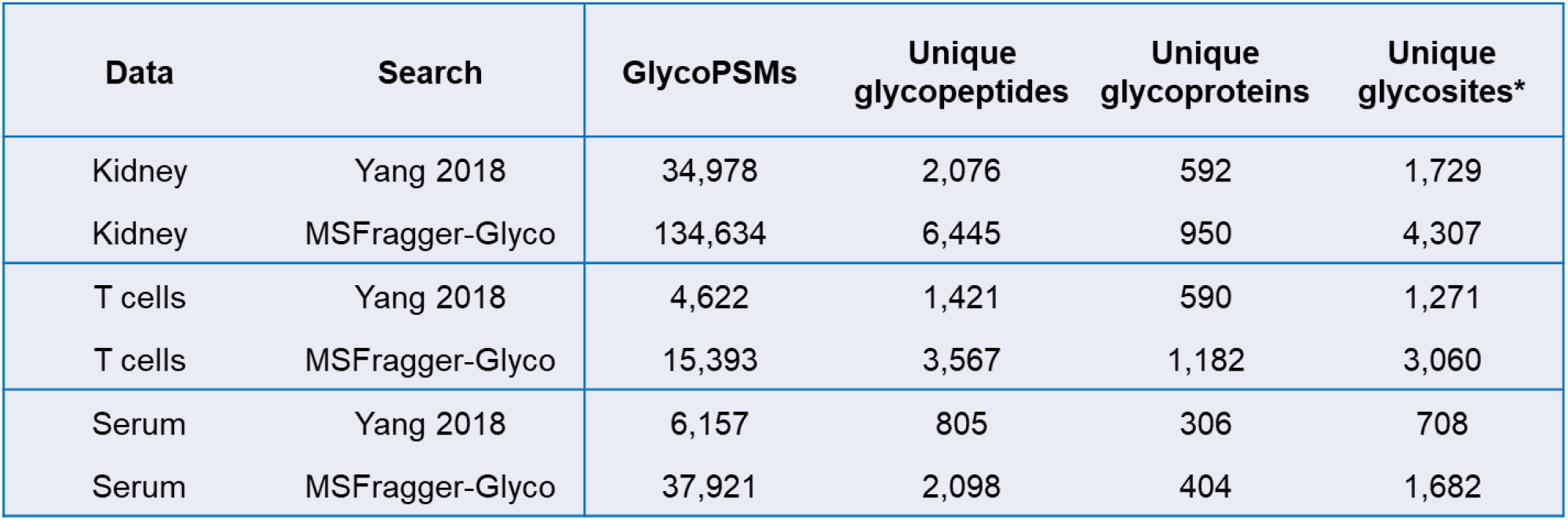
Comparison of MSFragger and Yang *et al*. results for all sample types. The MSFragger search is considering 110 possible glycan compositions as well as more missed cleavages and variable modifications than the Yang *et al*. search in each case. *Glycosites are computed as in Yang *et al*., assuming the peptide N-terminal Ser or Thr is the only glycosite in the peptide, for purposes of comparison. Our results indicate that many of these peptides in fact have multiple glycosylation sites, so the true number of glycosites is likely higher in both our searches and those of Yang *et al*.

**Figure 4.**
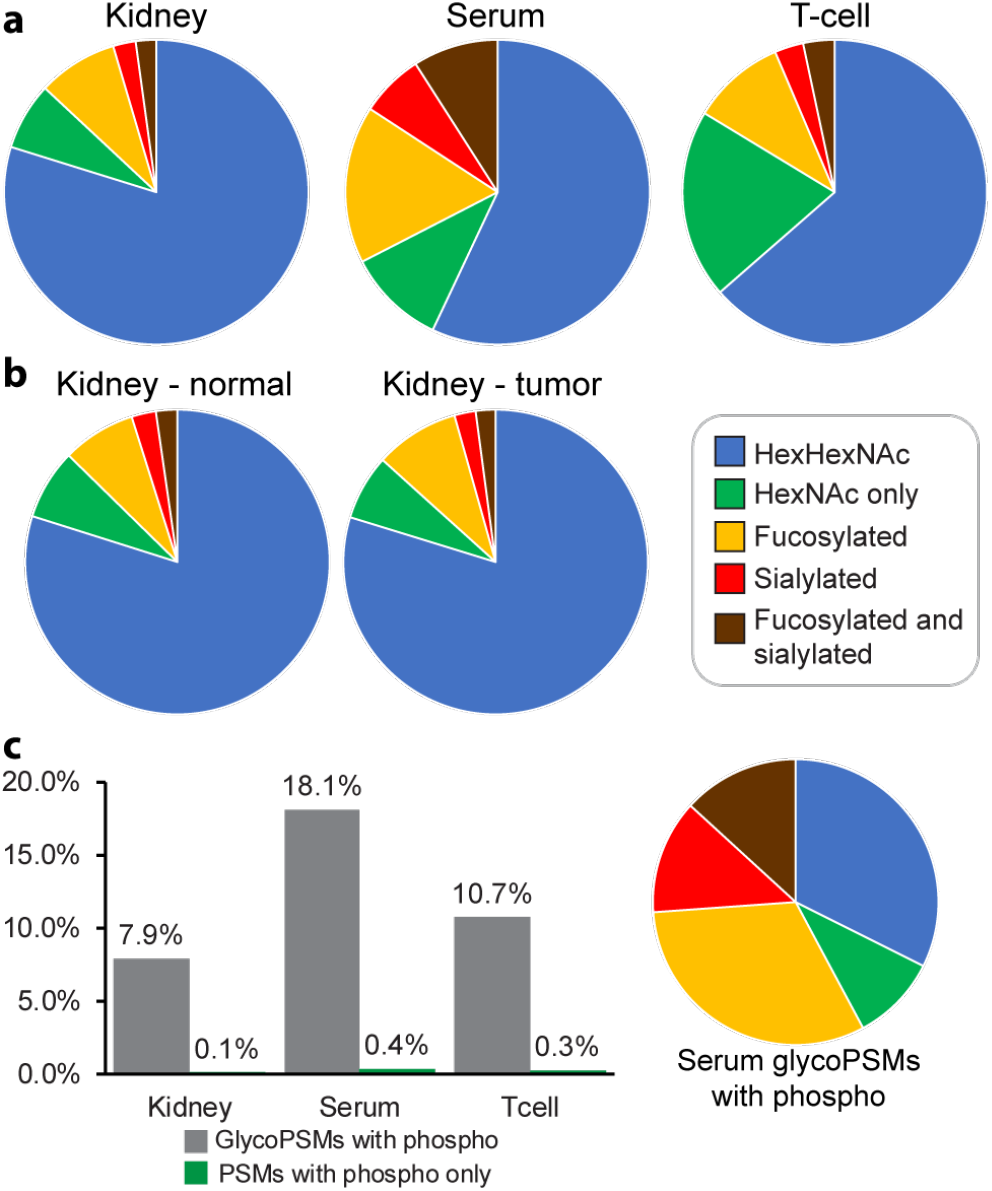
O-glycan composition types compared across tissues. a) Kidney (all), Serum, and T-cells compared. Note that HexHexNAc category may include some “HexNAc only” compositions if they coincide on the same peptide with a HexHexNAc glycan. Serum samples contained a much higher proportion of fucosylated and sialylated glycans than kidney or T-cell samples. b) Normal and tumor kidney tissue compositions compared, showing minimal difference in observed compositions. c) left, PSMs containing both glycosylation and phosphorylation shown as a percent of the total number of glycoPSMs for kidney, serum, and T-cell samples. Right, compositions of serum glycoPSMs also containing phosphorylation showing enrichment for fucosylation and sialylation relative to all glycoPSMs.

The ultrafast speed of MSFragger-Glyco, enabling both open and glycan mass offset searches, thus represents a powerful platform for delving deeply into complex O-glycoproteomics data. The mass offset method is particularly effective for O-glycoproteomics given the highly labile nature of O-glycosylation and vast numbers of potential compositions. The complexity of data from Yang *et al*., with many missed cleavages and several abundant variable modifications in addition to complex O-glycosylation, demonstrates the some of the challenges in comprehensively analyzing O-glycoproteomics data and the ability of MSFragger-Glyco to overcome these challenges.

## Discussion

Here, we demonstrate that MSFragger-Glyco provides superior performance for labile modifications, which we use to dramatically improve annotation of glycopeptide spectra. In reanalyzing several large-scale glycoproteomics datasets, we provide increases of 83-520% in the total number of glycoPSMs annotated, ultimately identifying many more glycoproteins and glycosylation sites from the same raw data. MSFragger-Glyco’s ultrafast index-based scoring enables complex searches for up to thousands of glycan compositions and several variable modifications, and even fully open searches, in reasonable time. We have demonstrated the utility of these capabilities for both N- and O-linked glycoproteomics data, revealing hundreds of additional glycosites in the N-linked data of Riley *et al*. and delving into the complexity of O-glycans found in several human samples from Yang *et al*., uncovering trends in composition and co-incidence with phosphorylation that were invisible to the original analysis.

While the mass offset search is substantially better than traditional methods at identifying glycopeptides from tandem MS data, and at localizing the site of a single labile glycan on a peptide, it offers mixed performance in localizing multiple glycans in a single peptide. The addition of *b/y* + HexNAc fragments provides additional information for localization over a variable modification search for labile glycans, offering potential for improved localization from HCD and hybrid activation data, particularly for N-glycans. However, multiple glycosylation sites on a single peptide are treated as a single mass offset, potentially resulting in poor localization if glycans are only partially fragmented. If there are sufficient glycan-containing fragment ions present, a post-search analysis could utilize the known peptide and glycans to perform a multi-site assignment. In the datasets we examined here, this issue was largely irrelevant as the coincidence of N-glycan sequons on a single peptide was rare and the energetic HCD fragmentation used in the O-glycan data resulted in nearly complete dissociation of all glycans, precluding direct localization.

As improvements to glycopeptide enrichment and analysis by MS continue, the quality and complexity of available data will continue to grow. MSFragger-Glyco provides a powerful platform to enable the next generation of glycoproteomics, combining a much-improved search method for labile glycans with the speed to perform very complex analyses. This method can be extended to glycoproteomic searches in other organisms and systems by changing to the glycan masses provided in the parameter file, or with simple changes to the code to include alternative potential glycosites. Similarly, MSFragger-Glyco can be used to search other labile modifications, such as ADP-ribosylation^46^, that are challenging to assess with conventional search strategies. These capabilities offer the potential to elucidate many areas of biology and disease that have proven challenging or even intractable, including the intricacies of O-glycosylation, crosstalk between glycosylation and other PTMs, and the many roles of glycosylation in disease.

## Acknowledgements

This work was funded in part by NIH grants R01-GM-094231 and U24-CA210967.

## Software and Data Availability

The program was developed in the cross-platform Java language and can be accessed at http://www.nesvilab.org/software

## Author Contributions

D.A.P., F.Y., and G.C.T. developed the algorithm. D.A.P. analyzed the data. A.I.N. conceived and supervised the project. D.A.P. and A.I.N. wrote the manuscript with input from all authors.

## Methods

Methods, including statements of data availability and any associated accession codes and references, are available in the online version of the paper.

### Data sets

N- and O-linked glycoproteomics data was downloaded from the PRIDE Archive^47^ with Proteome Xchange^48^ numbers PXD011533^10^ (N-glycan data) and PXD009476^12^ (O-glycan data). In PXD011533, N-glycopeptides from mouse brain tissue were lectin enriched (Concanavalin A) and analyzed by HCD-pd-AI-ETD LC-MS/MS on an Orbitrap Fusion Lumos mass spectrometer. Downloaded raw data files were centroided and converted to the mzML spectral format using MSConvert^49^, with HCD and AI-ETD scans filtered to separate mzML files. PXD008476 contains O-glycopeptides enriched from human kidney tissue, CEM T-cells, and human serum using an extraction procedure dubbed ExOO^12^. Enriched O-glycopeptides were analyzed by HCD LC-MS/MS on Q-Exactive HF Orbitrap mass spectrometer. Further details regarding sample preparation and MS analysis can be found in the corresponding publications.

### Open and mass offset searches with MSFragger-Glyco

The extension of open searching to include shifted ions for improved scoring with simultaneous localization of the mass shift (termed localization-aware open search) has been described elsewhere^22,31^. Briefly, searching “shifted ions,” or those resulting from the addition of a known (mass offset search) or unknown (open search) delta mass to a peptide sequence, as well as “regular ions,” or those resulting only from fragmentation of a database peptide, improves the sensitivity and quality of open (and mass offset) search results. Shifted ions can be indexed for rapid search by subtracting the delta mass from the observed precursor mass, enabling MSFragger to search shifted and regular fragment ions from a peptide simultaneously. Glycopeptide spectra are searched by providing potential glycan compositions as a list of allowed mass offsets. Raw spectra were deisotoped and de-charged in MSFragger-Glyco prior to analysis, which proved particularly helpful for high-mass glycopeptides. Spectra are searched in one of three modes, depending on the activation method (Fig. 1b). For CID/HCD (vibrational activation only), only regular ions are searched as the glycan is assumed to have dissociated from peptide (either entirely, or partially to form Y-ions or *b/y* ions with a single HexNAc remaining) (see Supplementary Table 1 for Y ions considered). For ETD/ECD (electronic activation only), regular and shifted ions are matched (as in a typical MSFragger open/mass offset search) assuming the glycan remains intact on the peptide. For hybrid activation (both vibrational and electronic), only regular *b/y*-type ions (no shifted *b/y*) are searched along with regular and shifted *c/z*-type ions to match all possible peptide and glycan fragmentation products simultaneously. A sequon check was added to ensure that only peptides with a potential glycosite are matched to a glycan mass offset, and an oxonium ion check to ensure that only spectra with evidence of glycan fragmentation can be matched to a glycopeptide (Supplementary Table 2). Y-type ions and a series of *b/y* ions plus the mass of a HexNAc were added to the fragment index for N-glycan mode (but not O-glycan).

### FDR control for glycoPSMs

Our FDR approach is designed for large-scale glycoproteomics, in which sufficiently many glycopeptide spectra are available for the target-decoy approach to FDR to be used, and is essentially the same as the procedure for FDR control of open search results. Filtering is performed with the Philosopher pipeline (https://philosopher.nesvilab.org/)^38^, including PeptideProphet modeling of peptide probabilities, ProteinProphet protein inference, and Philosopher’s internal filter for FDR control. A combined target and decoy (reversed) protein database is supplied to MSFragger-Glyco. Reversed N-glycan sequons are checked in reversed (decoy) peptides to ensure the same number of potential glycopeptides are searched in both target and decoy databases. The extended mass model of PeptideProphet is used as described in Kong *et al.^22^* to model probabilities for each mass offset (glycan mass). PeptideProphet parameters were as follows: extended mass model (4000 Da), glyc flag to separately model peptides containing the N-glycan sequon (for N-glycan data only), semi-parametric modeling using expectation scores only, cLevel −2. ProteinProphet was used with default settings except ‘maxppmdiff,’ which was set to 20,000,000 to ensure peptides containing glycan mass offsets were not filtered out. Philosopher filtering was performed at 1% PSM, peptide, and protein levels, followed by sequential filtering of PSMs from the final protein list.

The extended mass model of PeptideProphet functions as the primary method of controlling glycan-specific matching, though glycan FDR is not explicitly controlled. To validate that this appropriately controlled FDR for glycoPSMs, we performed several checks. First, FDR for PSMs containing each glycan offset all individually remain near 1% PSM FDR in all analyses performed (Supplementary Tables 7, 8). Second, searches were performed with an equal number of target and decoy glycans. Targets were the 10 most abundant glycans found in the data, and decoys were those glycan masses shifted by +20 Da (N-glycan) or +10 Da (O-glycan), after confirming there was no overlap with other known glycan compositions. In total, 0.6% (N-glycan) and 1.3% (O-glycan) of glycoPSMs were matched to decoy glycans (Supplementary Tables 3, 4), which broadly agree with the expected PSM FDR rate of 1%.

### Variable modification searches

A modified version of MSFragger was used to perform the comparative variable modification search for N-glycan data. 17 glycan masses were specified as variable modifications on N-X-S/T. The MSFragger code was modified to allow specification of the full sequon for a variable modification (standard MSFragger allows specification of single residues only). Oxidized Met was allowed (up to 2 per peptide) but no other variable modifications were allowed. Only 1 glycan was allowed per peptide. All other parameters were as in the N-glycan mass offset searches. For variable modification searches in O-glycan data, standard MSFragger was used with HexNAc-Hex (365.1322) and HexNAc (230.0897) specified on Ser/Thr residues (up to 3 each per peptide). Oxidation of Met (up to 2 per peptide) and guanidinylation of Lys (up to 2 per peptide) were allowed as well, and the maximum total number of variable modifications per peptide was set to 4. To match the search used in Yang *et al*., peptides of length 7 to 46 residues were considered, allowing up to 5 missed cleavages by OpeRATOR. All other parameters were identical to those used in the O-glycan mass offset searches.

### N-glycan mass offset search

MzML files were searched with MSFragger-Glyco using 182 mass offsets (Supplementary Table 5), identical to those used by Riley *et al*.^10^, against the glycoprotein database used by Riley *et al*. containing 3,574 entries with decoys added using Philosopher. Trypsin digestion with up to 3 missed cleavages was specified with variable modifications of oxidized Met, protein N-terminal acetylation, and peptide N-terminal pyroglutamate. Peptides containing the consensus sequon (N-X-S/T) and decoy (reversed) peptides containing the reversed sequon were considered as potential glycopeptides. Only spectra containing oxonium ion peaks with summed intensity at least 10% of the base peak were considered for glycan searches. Data was deisotoped and de-charged in MSFragger-Glyco, calibrated, and searched with mass tolerances for precursors and products of 20 and 10 ppm, respectively. Precursor and electron transfer-no dissociation peaks were removed, and data was square root-transformed prior to analysis. For AI-ETD data, *b,y,c,z*, and *Y* ions were considered in searching; for HCD data, *b,y,Y*and *b,y* + HexNAc ions were considered. No *b* or *y* ions containing the intact glycan were considered in either mode. Spectra were visualized using Byonic viewer and PDV^50^ to determine appropriate ion types during development. Search results from all raw files and both activation modes were processed together using Philosopher. PSMs and glycopeptides/proteins/sites were compared to those reported in the supporting information of Riley *et al*. and Liu *et al*.^8^ using custom Python scripts, pyOpenMS^51^, and Biopython^52^. Unique glycopeptides refer to unique peptide sequences present in at least one glycoPSM. Different glycans and/or other modifications to an existing sequence do not count as additional unique glycopeptides. Glycosites were assigned to the position of the glycan sequon (N-X-S/T) in each glycopeptide, with peptides containing multiple sites excluded from site-specific analyses.

### O-glycan mass offset search

Kidney, Serum, and T cell-derived samples were searched separately with MSFragger-Glyco using reviewed human sequences from UniProtKB (downloaded 08/22/2019, 20464 sequences in total) with decoys and common contaminants added using the Philosopher database command. The OpeRATOR enzyme used in Yang *et al.^12^* cleaves N-terminal to O-glycosylated Ser and Thr. Protein sequences were pre-digested at S/T with up to 10 missed cleavages, as not all Ser and Thr residues are glycosylated and the sites of glycosylation are not known in advance, except for 12-composition searches comparing directly to published results, in which 5 missed cleavages at Ser and Thr were allowed. The resulting peptides were introduced to MSFragger-Glyco as protein sequences in a custom database and digested with Trypsin, allowing 1 missed cleavage. Variable modifications of oxidation (M), guanidinylation and carbamidomethylation (K), phosphorylation (S/T/Y), and deamidation (Q) were specified after initial searches revealed high levels of each in the data. A list of 110 O-glycans (Supplementary Table 6) was curated from open search results on the data and passed to MSFragger as a mass offset list. Peptides were required to contain at least one S/T residue to be considered for glycan search, and spectra were required to have summed oxonium ion intensity at least 10% of the base peak. Data was deisotoped and de-charged in MSFragger-Glyco, calibrated with parameter optimization, and searched with mass tolerances for precursors and products of 20 ppm and 10 ppm, respectively. Precursor peaks were removed, and intensities were square root-transformed prior to analysis. Only unshifted (no glycan) *b* and *y* ions were considered in searching as very few spectra retained any glyco-related fragments following HCD. Filtering and validation were performed in Philosopher as for N-glycan AI-ETD data, with the exception of no glycan motif modeling in PeptideProphet.

