## Supporting information document for "Fast and Comprehensive N- and O-glycoproteomics analysis with MSFragger-Glyco"

### Table of Contents

1. **Supplementary Figure 1.** Example spectrum of a peptide containing multiple known sites of O-glycosylation.
2. **Supplementary Table 1.** Y ions used in MSFragger searches
3. **Supplementary Table 2.** Oxonium ions used in MSFragger searches
4. **Supplementary Table 3**. Results of decoy offset search for N-glycan data
5. **Supplementary Table 4**. Results of decoy offset search for O-glycan kidney data
6. **Supplementary Table 5**. 182 N-glycan compositions searched in Riley *et al.* data.
7. **Supplementary Table 6**. 110 O-glycan compositions searched in Yang *et al.* data.
8. **Supplementary Table 7**: FDR for each mass offset for N-glycan data from Riley *et al.*
9. **Supplementary Table 8**: FDR for each mass offset for O-glycan main search (110 glycans) from Yang *et al.*

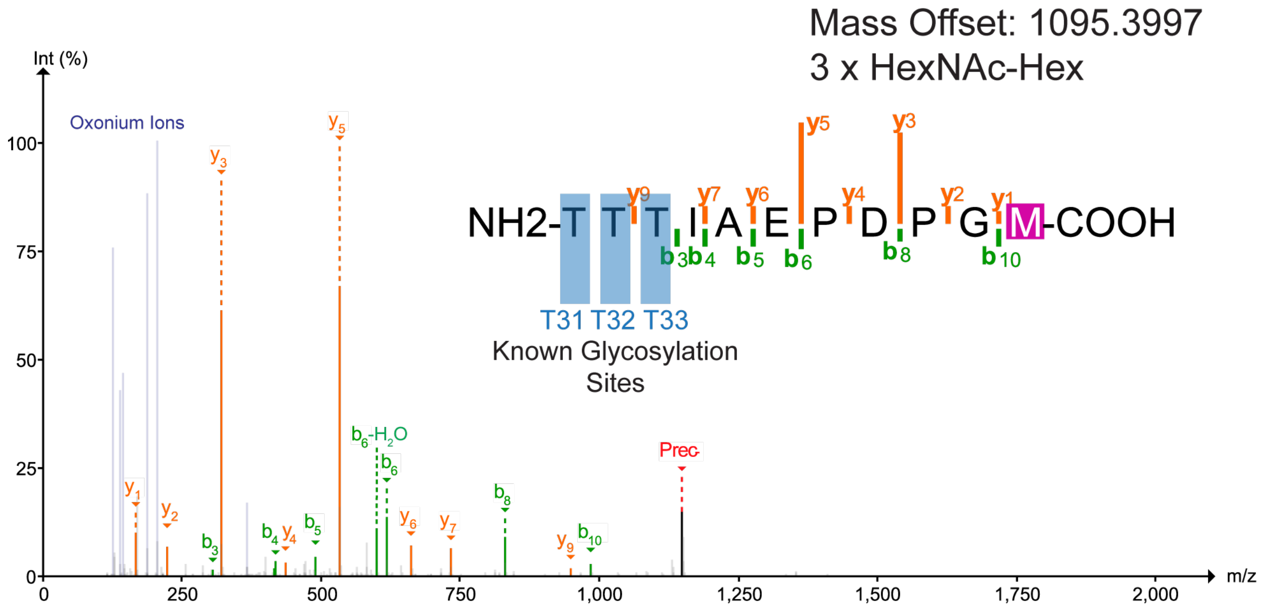


**Supplementary Figure 1.** Example spectrum of a peptide containing multiple known sites of O-glycosylation. No fragment ions bearing partial or intact glycan can be observed in the spectrum. All ions observed correspond to the fragmentation of the peptide sequence with no modifications aside from oxidation of Met-11. The mass offset of 1095.3997 corresponds to 3 copies of the common glycan HexNAcHex, which are likely present at the 3 Thr residues at the peptide N-terminus. All 3 Thr residues are known glycosylation sites of the protein (glycophorin-C).

**Supplementary Table 1.** Y ions used in MSFragger searches

| Mass | Composition |
|---|---|
| 203.07937 | HexNAc(1) |
| 406.15874 | HexNAc(2) |
| 568.21156 | HexNAc(2)Hex(1) |
| 730.26438 | HexNAc(2)Hex(2) |
| 892.3172 | HexNAc(2)Hex(3) |
| 1095.39657 | HexNAc(3)Hex(3) |
| 1257.44939 | HexNAc(3)Hex(4) |
| 349.137279 | HexNAc(1)Fuc(1) |
| 552.216649 | HexNAc(2)Fuc(1) |

**Supplementary Table 2.** Oxonium ions used in MSFragger searches

| Composition | Mass |
|---|---|
| HexNAc | 204.086646 |
| HexNAc - 18 | 186.076086 |
| HexNAcHex | 366.139466 |
| C6H10NO3 | 144.0656 |
| C7H8NO2 | 138.055 |
| C6H8NO2 | 126.055 |
| PhosphoHex | 243.026426 |
| Phospho2Hex | 405.079246 |
| 2Phospho2Hex | 485.045576 |
| Hex | 163.06 |

**Supplementary Table 3**. Results of decoy offset search for N-glycan data. The 10 most abundant mass offsets were searched along with 10 decoy offsets generated by adding 20 Da to each target mass offset. 0.6% of PSMs corresponded to decoy offsets, within the expected PSM FDR of 1%.

| Mass | Composition | PSMs | Decoy Mass | Decoy PSMs | FDR% |
|---|---|---|---|---|---|
| 892.3172 | HexNAc-2_Hex-3 | 926 | 912.3172 | 20 | 2.2% |
| 1038.3751 | HexNAc-2_Hex-3_Fuc-1 | 1417 | 1058.3751 | 21 | 1.5% |
| 1054.37 | HexNAc-2_Hex-4 | 2277 | 1074.37 | 11 | 0.5% |
| 1216.4228 | HexNAc-2_Hex-5 | 13485 | 1236.4228 | 45 | 0.3% |
| 1378.4757 | HexNAc-2_Hex-6 | 5999 | 1398.4757 | 19 | 0.3% |
| 1540.5285 | HexNAc-2_Hex-7 | 3846 | 1560.5285 | 8 | 0.2% |
| 1565.5601 | HexNAc-3_Hex-5_Fuc-1 | 392 | 1585.5601 | 9 | 2.3% |
| 1702.5813 | HexNAc-2_Hex-8 | 4140 | 1722.5813 | 50 | 1.2% |
| 1768.6395 | HexNAc-4_Hex-5_Fuc-1 | 602 | 1788.6395 | 17 | 2.8% |
| 1864.6341 | HexNAc-2_Hex-9 | 2048 | 1884.6341 | 14 | 0.7% |
|  | **Target Total:** | **35132** | **Decoy Total:** | **214** | **0.6%** |

**Supplementary Table 4**. Results of decoy offset search for O-glycan kidney data. The 10 most abundant mass offsets were searched along with 10 decoy offsets generated by adding 10 Da to each target mass offset. 10 Da was chosen to avoid conflicts with other known glycan composition masses and isotope distributions. In total, 1.3% of PSMs corresponded to decoy offsets, close to the expected PSM FDR of 1%.

| Mass | Composition | PSMs | Decoy Mass | Decoy PSMs | FDR% |
|---|---|---|---|---|---|
| 203.0794 | HexNAc-1 | 5425 | 213.0794 | 62 | 1.1% |
| 365.1322 | HexNAc-1_Hex-1 | 45900 | 375.1322 | 201 | 0.4% |
| 568.2116 | HexNAc-2_Hex-1 | 2783 | 578.2116 | 65 | 2.3% |
| 730.2644 | HexNAc-2_Hex-2 | 12322 | 740.2644 | 113 | 0.9% |
| 771.2909 | HexNAc-3_Hex-1 | 1172 | 781.2909 | 160 | 13.7% |
| 933.3438 | HexNAc-3_Hex-2 | 964 | 943.3438 | 163 | 16.9% |
| 1095.3966 | HexNAc-3_Hex-3 | 6379 | 1105.3966 | 113 | 1.8% |
| 1298.4759 | HexNAc-4_Hex-3 | 441 | 1308.4759 | 23 | 5.2% |
| 1460.5288 | HexNAc-4_Hex-4 | 1702 | 1470.5288 | 100 | 5.9% |
| 1825.6609 | HexNAc-5_Hex-5 | 885 | 1835.6609 | 17 | 1.9% |
| | **Target Total:** | **77973** | **Decoy Total:** | **1017** | **1.3%** |

**Supplementary Table 5**. 182 N-glycan compositions searched in Riley *et al.* data.

| Mass | Composition | Mass | Composition |
|---|---|---|---|
| 203.07937 | HexNAc(1) | 2715.962512 | HexNAc(5)Hex(6)Fuc(1)NeuAc(2) |
| 349.137279 | HexNAc(1)Fuc(1) | 3007.057929 | HexNAc(5)Hex(6)Fuc(1)NeuAc(3) |
| 406.15874 | HexNAc(2) | 2279.829588 | HexNAc(5)Hex(6)Fuc(2) |
| 552.216649 | HexNAc(2)Fuc(1) | 2570.925005 | HexNAc(5)Hex(6)Fuc(2)NeuAc(1) |
| 568.21156 | HexNAc(2)Hex(1) | 2425.887497 | HexNAc(5)Hex(6)Fuc(3) |
| 714.269469 | HexNAc(2)Hex(1)Fuc(1) | 2716.982914 | HexNAc(5)Hex(6)Fuc(3)NeuAc(1) |
| 2026.68694 | HexNAc(2)Hex(10) | 3008.07833 | HexNAc(5)Hex(6)Fuc(3)NeuAc(2) |
| 2188.73976 | HexNAc(2)Hex(11) | 3299.173747 | HexNAc(5)Hex(6)Fuc(3)NeuAc(3) |
| 2350.79258 | HexNAc(2)Hex(12) | 2571.945406 | HexNAc(5)Hex(6)Fuc(4) |
| 730.26438 | HexNAc(2)Hex(2) | 2863.040823 | HexNAc(5)Hex(6)Fuc(4)NeuAc(1) |
| 876.322289 | HexNAc(2)Hex(2)Fuc(1) | 2278.809187 | HexNAc(5)Hex(6)NeuAc(1) |
| 892.3172 | HexNAc(2)Hex(3) | 2569.904603 | HexNAc(5)Hex(6)NeuAc(2) |
| 1038.375109 | HexNAc(2)Hex(3)Fuc(1) | 2861.00002 | HexNAc(5)Hex(6)NeuAc(3) |
| 1054.37002 | HexNAc(2)Hex(4) | 2586.919916 | HexNAc(5)Hex(7)Fuc(1)NeuAc(1) |
| 1200.427929 | HexNAc(2)Hex(4)Fuc(1) | 2878.015332 | HexNAc(5)Hex(7)Fuc(1)NeuAc(2) |
| 1216.42284 | HexNAc(2)Hex(5) | 2311.81941 | HexNAc(5)Hex(8) |
| 1362.480749 | HexNAc(2)Hex(5)Fuc(1) | 2457.877319 | HexNAc(5)Hex(8)Fuc(1) |
| 1378.47566 | HexNAc(2)Hex(6) | 2619.930139 | HexNAc(5)Hex(9)Fuc(1) |
| 1524.533569 | HexNAc(2)Hex(6)Fuc(1) | 1704.63468 | HexNAc(6)Hex(3) |
| 1458.44199 | HexNAc(2)Hex(6)Phospho(1) | 1850.692589 | HexNAc(6)Hex(3)Fuc(1) |
| 1540.52848 | HexNAc(2)Hex(7) | 2141.788006 | HexNAc(6)Hex(3)Fuc(1)NeuAc(1) |
| 1686.586389 | HexNAc(2)Hex(7)Fuc(1) | 2432.883422 | HexNAc(6)Hex(3)Fuc(1)NeuAc(2) |
| 1702.5813 | HexNAc(2)Hex(8) | 1996.750498 | HexNAc(6)Hex(3)Fuc(2) |
| 1864.63412 | HexNAc(2)Hex(9) | 1866.6875 | HexNAc(6)Hex(4) |
| 1095.39657 | HexNAc(3)Hex(3) | 2012.745409 | HexNAc(6)Hex(4)Fuc(1) |
| 1241.454479 | HexNAc(3)Hex(3)Fuc(1) | 2158.803318 | HexNAc(6)Hex(4)Fuc(2) |
| 1257.44939 | HexNAc(3)Hex(4) | 2157.782917 | HexNAc(6)Hex(4)NeuAc(1) |
| 1403.507299 | HexNAc(3)Hex(4)Fuc(1) | 2028.74032 | HexNAc(6)Hex(5) |
| 1694.602716 | HexNAc(3)Hex(4)Fuc(1)NeuAc(1) | 2174.798229 | HexNAc(6)Hex(5)Fuc(1) |
| 1549.565208 | HexNAc(3)Hex(4)Fuc(2) | 2465.893646 | HexNAc(6)Hex(5)Fuc(1)NeuAc(1) |
| 1840.660625 | HexNAc(3)Hex(4)Fuc(2)NeuAc(1) | 2756.989062 | HexNAc(6)Hex(5)Fuc(1)NeuAc(2) |
| 1548.544807 | HexNAc(3)Hex(4)NeuAc(1) | 3048.084479 | HexNAc(6)Hex(5)Fuc(1)NeuAc(3) |
| 1419.50221 | HexNAc(3)Hex(5) | 2320.856138 | HexNAc(6)Hex(5)Fuc(2) |
| 1565.560119 | HexNAc(3)Hex(5)Fuc(1) | 2611.951555 | HexNAc(6)Hex(5)Fuc(2)NeuAc(1) |
| 1856.655536 | HexNAc(3)Hex(5)Fuc(1)NeuAc(1) | 2466.914047 | HexNAc(6)Hex(5)Fuc(3) |
| 2147.750952 | HexNAc(3)Hex(5)Fuc(1)NeuAc(2) | 2190.79314 | HexNAc(6)Hex(6) |
| 1710.597627 | HexNAc(3)Hex(5)NeuAc(1) | 2336.851049 | HexNAc(6)Hex(6)Fuc(1) |
| 1581.55503 | HexNAc(3)Hex(6) | 2627.946466 | HexNAc(6)Hex(6)Fuc(1)NeuAc(1) |
| 1727.612939 | HexNAc(3)Hex(6)Fuc(1) | 2482.908958 | HexNAc(6)Hex(6)Fuc(2) |
| 2018.708356 | HexNAc(3)Hex(6)Fuc(1)NeuAc(1) | 2774.004375 | HexNAc(6)Hex(6)Fuc(2)NeuAc(1) |
| 1872.650447 | HexNAc(3)Hex(6)NeuAc(1) | 3065.099791 | HexNAc(6)Hex(6)Fuc(2)NeuAc(2) |
| 1298.47594 | HexNAc(4)Hex(3) | 2481.888557 | HexNAc(6)Hex(6)NeuAc(1) |
| 1444.533849 | HexNAc(4)Hex(3)Fuc(1) | 2772.983973 | HexNAc(6)Hex(6)NeuAc(2) |
| 1589.571357 | HexNAc(4)Hex(3)NeuAc(1) | 3064.07939 | HexNAc(6)Hex(6)NeuAc(3) |

| | | | |
|---|---|---|---|
| 1460.52876 | HexNAc(4)Hex(4) | 2352.84596 | HexNAc(6)Hex(7) |
| 1606.586669 | HexNAc(4)Hex(4)Fuc(1) | 2789.999286 | HexNAc(6)Hex(7)Fuc(1)NeuAc(1) |
| 1897.682086 | HexNAc(4)Hex(4)Fuc(1)NeuAc(1) | 3081.094702 | HexNAc(6)Hex(7)Fuc(1)NeuAc(2) |
| 1752.644578 | HexNAc(4)Hex(4)Fuc(2) | 2644.961778 | HexNAc(6)Hex(7)Fuc(2) |
| 1751.624177 | HexNAc(4)Hex(4)NeuAc(1) | 2936.057195 | HexNAc(6)Hex(7)Fuc(2)NeuAc(1) |
| 1622.58158 | HexNAc(4)Hex(5) | 2791.019687 | HexNAc(6)Hex(7)Fuc(3) |
| 1768.639489 | HexNAc(4)Hex(5)Fuc(1) | 3082.115104 | HexNAc(6)Hex(7)Fuc(3)NeuAc(1) |
| 2059.734906 | HexNAc(4)Hex(5)Fuc(1)NeuAc(1) | 2643.941377 | HexNAc(6)Hex(7)NeuAc(1) |
| 2350.830322 | HexNAc(4)Hex(5)Fuc(1)NeuAc(2) | 2935.036793 | HexNAc(6)Hex(7)NeuAc(2) |
| 1914.697398 | HexNAc(4)Hex(5)Fuc(2) | 3226.13221 | HexNAc(6)Hex(7)NeuAc(3) |
| 2205.792815 | HexNAc(4)Hex(5)Fuc(2)NeuAc(1) | 3517.227626 | HexNAc(6)Hex(7)NeuAc(4) |
| 2496.888231 | HexNAc(4)Hex(5)Fuc(2)NeuAc(2) | 2952.052106 | HexNAc(6)Hex(8)Fuc(1)NeuAc(1) |
| 2351.850724 | HexNAc(4)Hex(5)Fuc(3)NeuAc(1) | 2805.994197 | HexNAc(6)Hex(8)NeuAc(1) |
| 2642.94614 | HexNAc(4)Hex(5)Fuc(3)NeuAc(2) | 2676.9516 | HexNAc(6)Hex(9) |
| 1913.676997 | HexNAc(4)Hex(5)NeuAc(1) | 3114.104926 | HexNAc(6)Hex(9)Fuc(1)NeuAc(1) |
| 2204.772413 | HexNAc(4)Hex(5)NeuAc(2) | 3405.200342 | HexNAc(6)Hex(9)Fuc(1)NeuAc(2) |
| 1784.6344 | HexNAc(4)Hex(6) | 1907.71405 | HexNAc(7)Hex(3) |
| 1930.692309 | HexNAc(4)Hex(6)Fuc(1) | 2053.771959 | HexNAc(7)Hex(3)Fuc(1) |
| 2221.787726 | HexNAc(4)Hex(6)Fuc(1)NeuAc(1) | 2069.76687 | HexNAc(7)Hex(4) |
| 2076.750218 | HexNAc(4)Hex(6)Fuc(2) | 2215.824779 | HexNAc(7)Hex(4)Fuc(1) |
| 2075.729817 | HexNAc(4)Hex(6)NeuAc(1) | 2393.87251 | HexNAc(7)Hex(6) |
| 1946.68722 | HexNAc(4)Hex(7) | 2539.930419 | HexNAc(7)Hex(6)Fuc(1) |
| 2092.745129 | HexNAc(4)Hex(7)Fuc(1) | 2555.92533 | HexNAc(7)Hex(7) |
| 2237.782637 | HexNAc(4)Hex(7)NeuAc(1) | 2701.983239 | HexNAc(7)Hex(7)Fuc(1) |
| 1501.55531 | HexNAc(5)Hex(3) | 3575.269489 | HexNAc(7)Hex(7)Fuc(1)NeuAc(3) |
| 1647.613219 | HexNAc(5)Hex(3)Fuc(1) | 2717.97815 | HexNAc(7)Hex(8) |
| 1938.708636 | HexNAc(5)Hex(3)Fuc(1)NeuAc(1) | 2864.036059 | HexNAc(7)Hex(8)Fuc(1) |
| 1793.671128 | HexNAc(5)Hex(3)Fuc(2) | 3155.131476 | HexNAc(7)Hex(8)Fuc(1)NeuAc(1) |
| 1663.60813 | HexNAc(5)Hex(4) | 4028.417725 | HexNAc(7)Hex(8)Fuc(1)NeuAc(4) |
| 1809.666039 | HexNAc(5)Hex(4)Fuc(1) | 3009.073567 | HexNAc(7)Hex(8)NeuAc(1) |
| 2100.761456 | HexNAc(5)Hex(4)Fuc(1)NeuAc(1) | 2110.79342 | HexNAc(8)Hex(3) |
| 2391.856872 | HexNAc(5)Hex(4)Fuc(1)NeuAc(2) | 2256.851329 | HexNAc(8)Hex(3)Fuc(1) |
| 1955.723948 | HexNAc(5)Hex(4)Fuc(2) | 2272.84624 | HexNAc(8)Hex(4) |
| 1954.703547 | HexNAc(5)Hex(4)NeuAc(1) | 2434.89906 | HexNAc(8)Hex(5) |
| 2245.798963 | HexNAc(5)Hex(4)NeuAc(2) | 2580.956969 | HexNAc(8)Hex(5)Fuc(1) |
| 1825.66095 | HexNAc(5)Hex(5) | 2759.0047 | HexNAc(8)Hex(7) |
| 1971.718859 | HexNAc(5)Hex(5)Fuc(1) | 2921.05752 | HexNAc(8)Hex(8) |
| 2262.814276 | HexNAc(5)Hex(5)Fuc(1)NeuAc(1) | 3083.11034 | HexNAc(8)Hex(9) |
| 2553.909692 | HexNAc(5)Hex(5)Fuc(1)NeuAc(2) | 3229.168249 | HexNAc(8)Hex(9)Fuc(1) |
| 2117.776768 | HexNAc(5)Hex(5)Fuc(2) | 3448.24253 | HexNAc(9)Hex(10) |
| 2408.872185 | HexNAc(5)Hex(5)Fuc(2)NeuAc(1) | 2313.87279 | HexNAc(9)Hex(3) |
| 2263.834677 | HexNAc(5)Hex(5)Fuc(3) | 2459.930699 | HexNAc(9)Hex(3)Fuc(1) |
| 2116.756367 | HexNAc(5)Hex(5)NeuAc(1) | 2475.92561 | HexNAc(9)Hex(4) |
| 2407.851783 | HexNAc(5)Hex(5)NeuAc(2) | 2621.983519 | HexNAc(9)Hex(4)Fuc(1) |
| 1987.71377 | HexNAc(5)Hex(6) | 2800.03125 | HexNAc(9)Hex(6) |
| 2133.771679 | HexNAc(5)Hex(6)Fuc(1) | 2946.089159 | HexNAc(9)Hex(6)Fuc(1) |
| 2424.867096 | HexNAc(5)Hex(6)Fuc(1)NeuAc(1) | 3432.247619 | HexNAc(9)Hex(9)Fuc(1) |

**Supplementary Table 6**. 110 O-glycan compositions searched in Yang *et al.* data.

| Mass | Composition | Mass | Composition |
|---|---|---|---|
| 146.0579 | Fuc(1) | 1257.4494 | HexNAc(3)Hex(4) |
| 203.0794 | HexNAc(1) | 1282.481 | HexNAc(4)Hex(2)Fuc(1) |
| 349.1373 | HexNAc(1)Fuc(1) | 1298.4759 | HexNAc(4)Hex(3) |
| 365.1322 | HexNAc(1)Hex(1) | 1307.5127 | HexNAc(5)Fuc(2) |
| 406.1587 | HexNAc(2) | 1321.4542 | HexNAc(1)Hex(6)Fuc(1) |
| 494.1748 | HexNAc(1)NeuAc(1) | 1339.5025 | HexNAc(5)Hex(2) |
| 511.1901 | HexNAc(1)Hex(1)Fuc(1) | 1354.5021 | HexNAc(3)Hex(1)Fuc(2)NeuAc(1) |
| 527.185 | HexNAc(1)Hex(2) | 1362.4807 | HexNAc(2)Hex(5)Fuc(1) |
| 568.2116 | HexNAc(2)Hex(1) | 1378.4757 | HexNAc(2)Hex(6) |
| 609.2381 | HexNAc(3) | 1386.492 | HexNAc(3)Hex(3)NeuAc(1) |
| 640.2327 | HexNAc(1)Fuc(1)NeuAc(1) | 1395.5287 | HexNAc(4)Fuc(2)NeuAc(1) |
| 656.2276 | HexNAc(1)Hex(1)NeuAc(1) | 1403.5073 | HexNAc(3)Hex(4)Fuc(1) |
| 673.2429 | HexNAc(1)Hex(2)Fuc(1) | 1444.5338 | HexNAc(4)Hex(3)Fuc(1) |
| 689.2378 | HexNAc(1)Hex(3) | 1460.5288 | HexNAc(4)Hex(4) |
| 697.2541 | HexNAc(2)NeuAc(1) | 1501.5553 | HexNAc(5)Hex(3) |
| 698.2746 | HexNAc(2)Fuc(2) | 1524.5336 | HexNAc(2)Hex(6)Fuc(1) |
| 714.2695 | HexNAc(2)Hex(1)Fuc(1) | 1532.5499 | HexNAc(3)Hex(3)Fuc(1)NeuAc(1) |
| 730.2644 | HexNAc(2)Hex(2) | 1540.5285 | HexNAc(2)Hex(7) |
| 755.296 | HexNAc(3)Fuc(1) | 1548.5448 | HexNAc(3)Hex(4)NeuAc(1) |
| 771.2909 | HexNAc(3)Hex(1) | 1557.5815 | HexNAc(4)Hex(1)Fuc(2)NeuAc(1) |
| 786.2906 | HexNAc(1)Fuc(2)NeuAc(1) | 1565.5601 | HexNAc(3)Hex(5)Fuc(1) |
| 802.2855 | HexNAc(1)Hex(1)Fuc(1)NeuAc(1) | 1573.5764 | HexNAc(4)Hex(2)Fuc(1)NeuAc(1) |
| 812.3175 | HexNAc(4) | 1574.5968 | HexNAc(4)Hex(2)Fuc(3) |
| 818.2804 | HexNAc(1)Hex(2)NeuAc(1) | 1589.5713 | HexNAc(4)Hex(3)NeuAc(1) |
| 835.2957 | HexNAc(1)Hex(3)Fuc(1) | 1606.5867 | HexNAc(4)Hex(4)Fuc(1) |
| 843.312 | HexNAc(2)Fuc(1)NeuAc(1) | 1622.5816 | HexNAc(4)Hex(5) |
| 851.2907 | HexNAc(1)Hex(4) | 1647.6132 | HexNAc(5)Hex(3)Fuc(1) |
| 859.307 | HexNAc(2)Hex(1)NeuAc(1) | 1663.6081 | HexNAc(5)Hex(4) |
| 876.3223 | HexNAc(2)Hex(2)Fuc(1) | 1686.5864 | HexNAc(2)Hex(7)Fuc(1) |
| 892.3172 | HexNAc(2)Hex(3) | 1704.6347 | HexNAc(6)Hex(3) |
| 917.3488 | HexNAc(3)Hex(1)Fuc(1) | 1727.6129 | HexNAc(3)Hex(6)Fuc(1) |
| 933.3438 | HexNAc(3)Hex(2) | 1752.6446 | HexNAc(4)Hex(4)Fuc(2) |
| 974.3703 | HexNAc(4)Hex(1) | 1768.6395 | HexNAc(4)Hex(5)Fuc(1) |
| 989.37 | HexNAc(2)Fuc(2)NeuAc(1) | 1825.6609 | HexNAc(5)Hex(5) |
| 997.3486 | HexNAc(1)Hex(4)Fuc(1) | 1831.6239 | HexNAc(2)Hex(7)NeuAc(1) |
| 1005.3649 | HexNAc(2)Hex(1)Fuc(1)NeuAc(1) | 1866.6875 | HexNAc(6)Hex(4) |
| 1006.3853 | HexNAc(2)Hex(1)Fuc(3) | 1905.6607 | HexNAc(3)Hex(8) |
| 1013.3435 | HexNAc(1)Hex(5) | 1913.677 | HexNAc(4)Hex(5)NeuAc(1) |
| 1015.3969 | HexNAc(5) | 1914.6974 | HexNAc(4)Hex(5)Fuc(2) |
| 1021.3598 | HexNAc(2)Hex(2)NeuAc(1) | 1930.6923 | HexNAc(4)Hex(6)Fuc(1) |
| 1038.3751 | HexNAc(2)Hex(3)Fuc(1) | 1971.7189 | HexNAc(5)Hex(5)Fuc(1) |

| | | | |
|---|---|---|---|
| 1046.3914 | HexNAc(3)Fuc(1)NeuAc(1) | 1987.7138 | HexNAc(5)Hex(6) |
| 1054.37 | HexNAc(2)Hex(4) | 2018.7083 | HexNAc(3)Hex(6)Fuc(1)NeuAc(1) |
| 1079.4017 | HexNAc(3)Hex(2)Fuc(1) | 2028.7403 | HexNAc(6)Hex(5) |
| 1095.3966 | HexNAc(3)Hex(3) | 2092.7451 | HexNAc(4)Hex(7)Fuc(1) |
| 1136.4231 | HexNAc(4)Hex(2) | 2117.7768 | HexNAc(5)Hex(5)Fuc(2) |
| 1151.4228 | HexNAc(2)Hex(1)Fuc(2)NeuAc(1) | 2133.7717 | HexNAc(5)Hex(6)Fuc(1) |
| 1159.4014 | HexNAc(1)Hex(5)Fuc(1) | 2174.7982 | HexNAc(6)Hex(5)Fuc(1) |
| 1183.4126 | HexNAc(2)Hex(3)NeuAc(1) | 2190.7931 | HexNAc(6)Hex(6) |
| 1192.4493 | HexNAc(3)Fuc(2)NeuAc(1) | 2213.7714 | HexNAc(3)Hex(9)Fuc(1) |
| 1200.4279 | HexNAc(2)Hex(4)Fuc(1) | 2352.846 | HexNAc(6)Hex(7) |
| 1216.4228 | HexNAc(2)Hex(5) | 2393.8725 | HexNAc(7)Hex(6) |
| 1218.4762 | HexNAc(6) | 2498.9039 | HexNAc(6)Hex(7)Fuc(1) |
| 1224.4391 | HexNAc(3)Hex(2)NeuAc(1) | 2555.9253 | HexNAc(7)Hex(7) |
| 1241.4545 | HexNAc(3)Hex(3)Fuc(1) | 2701.9832 | HexNAc(7)Hex(7)Fuc(1) |

**Supplementary Table 7**. FDR for each mass offset for N-glycan data from Riley *et al.*

| Modification Mass | Target PSMs | Decoy PSMs | **FDR %** | Modification Mass | Target PSMs | Decoy PSMs | **FDR %** |
|---|---|---|---|---|---|---|---|
| 0 | 54599 | 11 | **0** | 2864 | 5 | 0 | **0** |
| 16 | 42657 | 11 | **0** | 2466.9 | 5 | 0 | **0** |
| 1216.4 | 9516 | 17 | **0.2** | 1851.7 | 5 | 0 | **0** |
| 1378.5 | 4297 | 13 | **0.3** | 1987.8 | 5 | 0 | **0** |
| 1217.4 | 3312 | 7 | **0.2** | 2223.8 | 5 | 1 | **20** |
| 1702.6 | 2773 | 3 | **0.1** | 2217.9 | 5 | 0 | **0** |
| 1540.5 | 2627 | 0 | **0** | 2459.9 | 5 | 0 | **0** |
| 1054.4 | 1810 | 9 | **0.5** | 2947.1 | 5 | 0 | **0** |
| 1379.5 | 1466 | 2 | **0.1** | 3067.1 | 5 | 0 | **0** |
| 1864.6 | 1310 | 0 | **0** | 2013.7 | 5 | 0 | **0** |
| 1703.6 | 1215 | 1 | **0.1** | 2679 | 4 | 0 | **0** |
| 1038.4 | 1144 | 0 | **0** | 3576.3 | 4 | 0 | **0** |
| 1541.5 | 1081 | 2 | **0.2** | 2936 | 4 | 0 | **0** |
| 1218.4 | 929 | 4 | **0.4** | 1526.5 | 4 | 0 | **0** |
| 892.3 | 807 | 5 | **0.6** | 2436.9 | 4 | 0 | **0** |
| 1865.7 | 720 | 1 | **0.1** | 1696.6 | 4 | 0 | **0** |
| 1704.6 | 440 | 0 | **0** | 2569.9 | 4 | 0 | **0** |
| 1380.5 | 432 | 1 | **0.2** | 3085.1 | 4 | 0 | **0** |
| 1055.4 | 420 | 0 | **0** | 2281.9 | 4 | 0 | **0** |
| 42 | 420 | 0 | **0** | 1608.6 | 4 | 0 | **0** |
| 1768.7 | 403 | 0 | **0** | 2191.9 | 4 | 0 | **0** |
| 1241.5 | 389 | 1 | **0.3** | 2407.8 | 4 | 0 | **0** |
| -17 | 350 | 2 | **0.6** | 3432.2 | 4 | 0 | **0** |
| 1458.5 | 339 | 1 | **0.3** | 1665.6 | 4 | 0 | **0** |
| 1542.6 | 336 | 0 | **0** | 2581.9 | 4 | 0 | **0** |
| 1565.6 | 276 | 2 | **0.7** | 570.2 | 4 | 0 | **0** |
| 1362.5 | 249 | 2 | **0.8** | 2257.8 | 4 | 0 | **0** |
| 1419.5 | 243 | 0 | **0** | 2238.8 | 4 | 0 | **0** |
| 1622.6 | 241 | 0 | **0** | 1794.7 | 4 | 0 | **0** |
| 1727.6 | 240 | 1 | **0.4** | 2110.7 | 4 | 0 | **0** |
| 1039.4 | 228 | 0 | **0** | 3227.1 | 4 | 0 | **0** |
| 730.3 | 226 | 0 | **0** | 1916.7 | 3 | 0 | **0** |
| 1873.7 | 212 | 0 | **0** | 1591.5 | 3 | 0 | **0** |
| 1866.6 | 207 | 0 | **0** | 2030.8 | 3 | 0 | **0** |
| 1930.7 | 192 | 1 | **0.5** | 2118.8 | 3 | 0 | **0** |
| 2018.7 | 170 | 0 | **0** | 2257 | 3 | 0 | **0** |
| 1459.5 | 162 | 0 | **0** | 2427.9 | 3 | 0 | **0** |
| 203.1 | 153 | 0 | **0** | 2761 | 3 | 0 | **0** |
| 893.3 | 149 | 0 | **0** | 1825.7 | 3 | 0 | **0** |
| 2433.9 | 138 | 0 | **0** | 2012.7 | 3 | 0 | **0** |

| | | | | | | | |
|---|---|---|---|---|---|---|---|
| 1769.7 | 136 | 0 | **0** | 2541 | 3 | 0 | **0** |
| 1444.5 | 126 | 0 | **0** | 3517.3 | 3 | 0 | **0** |
| 2076.8 | 117 | 1 | **0.9** | 2192.7 | 3 | 0 | **0** |
| 1874.7 | 115 | 0 | **0** | 2149.7 | 3 | 0 | **0** |
| 1566.6 | 114 | 0 | **0** | 2612 | 3 | 0 | **0** |
| 1897.7 | 113 | 0 | **0** | 2644 | 3 | 0 | **0** |
| 1420.5 | 109 | 0 | **0** | 2141.7 | 3 | 0 | **0** |
| 1872.7 | 105 | 0 | **0** | 2094.8 | 3 | 0 | **0** |
| 2432.9 | 104 | 0 | **0** | 3229.2 | 3 | 0 | **0** |
| 1623.6 | 103 | 0 | **0** | 2613.9 | 3 | 0 | **0** |
| 1931.7 | 101 | 0 | **0** | 2410.8 | 3 | 0 | **0** |
| 2019.7 | 97 | 0 | **0** | 1811.6 | 3 | 0 | **0** |
| 568.2 | 97 | 0 | **0** | 3066 | 3 | 0 | **0** |
| 1056.4 | 88 | 0 | **0** | 1957.7 | 3 | 0 | **0** |
| 2077.8 | 84 | 0 | **0** | 1947.7 | 3 | 0 | **0** |
| 1728.6 | 81 | 0 | **0** | 715.3 | 3 | 0 | **0** |
| 1752.7 | 81 | 0 | **0** | 1097.5 | 3 | 0 | **0** |
| 1242.5 | 80 | 0 | **0** | 2245.8 | 3 | 0 | **0** |
| 1705.6 | 77 | 0 | **0** | 2800 | 3 | 0 | **0** |
| 1040.4 | 76 | 0 | **0** | 2029.7 | 3 | 0 | **0** |
| 1363.5 | 76 | 0 | **0** | 3010.1 | 3 | 0 | **0** |
| 1200.4 | 75 | 0 | **0** | 2935 | 3 | 0 | **0** |
| 1581.6 | 74 | 0 | **0** | 2190.9 | 3 | 0 | **0** |
| 2434.9 | 68 | 0 | **0** | 2807.9 | 3 | 0 | **0** |
| 1095.4 | 66 | 0 | **0** | 3300.3 | 3 | 0 | **0** |
| 1257.5 | 61 | 0 | **0** | 2111.8 | 2 | 0 | **0** |
| 1784.7 | 55 | 0 | **0** | 1988.7 | 2 | 0 | **0** |
| 1445.5 | 53 | 0 | **0** | 2553.9 | 2 | 0 | **0** |
| 1460.5 | 52 | 0 | **0** | 2147.7 | 2 | 0 | **0** |
| 406.2 | 52 | 0 | **0** | 1711.6 | 2 | 0 | **0** |
| 2758 | 52 | 0 | **0** | 2716 | 2 | 0 | **0** |
| 2027.7 | 51 | 0 | **0** | 2677 | 2 | 0 | **0** |
| 1403.5 | 50 | 0 | **0** | 1590.6 | 2 | 0 | **0** |
| 1898.7 | 48 | 0 | **0** | 1842.7 | 2 | 0 | **0** |
| 1867.7 | 47 | 0 | **0** | 2720 | 2 | 0 | **0** |
| 1770.7 | 45 | 0 | **0** | 2938 | 2 | 0 | **0** |
| 2026.7 | 44 | 0 | **0** | 2702 | 2 | 0 | **0** |
| 1606.6 | 42 | 0 | **0** | 1827.7 | 2 | 0 | **0** |
| 1932.7 | 41 | 0 | **0** | 2542 | 2 | 0 | **0** |
| 731.3 | 40 | 0 | **0** | 2258.8 | 2 | 0 | **0** |
| 2757 | 38 | 0 | **0** | 1686.6 | 2 | 0 | **0** |
| 876.3 | 38 | 0 | **0** | 3228.1 | 2 | 0 | **0** |
| 1729.6 | 37 | 0 | **0** | 349.1 | 2 | 0 | **0** |

| | | | | | | | |
|---|---|---|---|---|---|---|---|
| 2020.7 | 36 | 0 | **0** | 2160.8 | 2 | 0 | **0** |
| 1582.6 | 35 | 0 | **0** | 2143.8 | 2 | 0 | **0** |
| 1201.4 | 35 | 0 | **0** | 1857.7 | 2 | 0 | **0** |
| 1421.5 | 33 | 0 | **0** | 2392.9 | 2 | 0 | **0** |
| 1753.7 | 33 | 0 | **0** | 3231.1 | 2 | 0 | **0** |
| 1624.6 | 33 | 0 | **0** | 2206.7 | 2 | 0 | **0** |
| 1785.7 | 33 | 0 | **0** | 2321.8 | 2 | 0 | **0** |
| 894.3 | 32 | 0 | **0** | 2496.9 | 2 | 0 | **0** |
| 2078.8 | 32 | 0 | **0** | 878.3 | 2 | 0 | **0** |
| 1243.5 | 31 | 0 | **0** | 3155.1 | 2 | 0 | **0** |
| 1647.6 | 30 | 0 | **0** | 552.2 | 2 | 0 | **0** |
| 1567.6 | 29 | 0 | **0** | 2354.9 | 2 | 0 | **0** |
| 204.1 | 29 | 0 | **0** | 2623.9 | 2 | 0 | **0** |
| 2435.9 | 29 | 0 | **0** | 2117.7 | 2 | 0 | **0** |
| 3082.1 | 28 | 0 | **0** | 2952 | 2 | 0 | **0** |
| 569.2 | 28 | 0 | **0** | 2801 | 2 | 0 | **0** |
| 2865.1 | 27 | 0 | **0** | 2425.9 | 2 | 0 | **0** |
| 1786.6 | 26 | 0 | **0** | 2189.7 | 2 | 0 | **0** |
| 2221.8 | 25 | 0 | **0** | 3115.1 | 2 | 0 | **0** |
| 732.3 | 25 | 0 | **0** | 2588.9 | 2 | 0 | **0** |
| 2921 | 24 | 0 | **0** | 1664.6 | 2 | 0 | **0** |
| 1549.6 | 22 | 0 | **0** | 3226.1 | 2 | 0 | **0** |
| 2759 | 22 | 0 | **0** | 2937 | 2 | 0 | **0** |
| 3083.1 | 22 | 0 | **0** | 1971.8 | 2 | 0 | **0** |
| 1364.5 | 21 | 0 | **0** | 2205.8 | 2 | 0 | **0** |
| 1648.6 | 21 | 0 | **0** | 1996.8 | 2 | 0 | **0** |
| 1259.4 | 20 | 0 | **0** | 3434.2 | 2 | 0 | **0** |
| 1298.5 | 20 | 0 | **0** | 2719 | 2 | 0 | **0** |
| 2866.1 | 20 | 0 | **0** | 1300.5 | 2 | 0 | **0** |
| 1649.6 | 19 | 0 | **0** | 2014.8 | 2 | 0 | **0** |
| 1938.7 | 18 | 0 | **0** | 1989.7 | 2 | 0 | **0** |
| 2222.8 | 18 | 0 | **0** | 2158.8 | 2 | 0 | **0** |
| 2946.1 | 18 | 0 | **0** | 2133.7 | 2 | 0 | **0** |
| 1913.7 | 18 | 0 | **0** | 2481.9 | 2 | 0 | **0** |
| 1460.4 | 17 | 0 | **0** | 1850.6 | 2 | 0 | **0** |
| 1751.6 | 17 | 0 | **0** | 2395 | 2 | 0 | **0** |
| 2351.8 | 17 | 1 | **5.9** | 2772.9 | 2 | 0 | **0** |
| 205.1 | 16 | 0 | **0** | 3114.2 | 1 | 0 | **0** |
| 1607.5 | 15 | 0 | **0** | 3065 | 1 | 0 | **0** |
| 2100.8 | 15 | 0 | **0** | 1840.6 | 1 | 0 | **0** |
| 1706.6 | 15 | 0 | **0** | 2054.7 | 1 | 0 | **0** |
| 1694.6 | 15 | 0 | **0** | 2279.8 | 1 | 0 | **0** |
| 1258.5 | 15 | 0 | **0** | 3448.3 | 1 | 0 | **0** |

| | | | | | | | |
|---|---|---|---|---|---|---|---|
| 1899.7 | 14 | 0 | **0** | 2313.8 | 1 | 0 | **0** |
| 2060.8 | 14 | 0 | **0** | 2393.8 | 1 | 0 | **0** |
| 2580.9 | 14 | 0 | **0** | 2702.9 | 1 | 0 | **0** |
| 1525.5 | 14 | 0 | **0** | 2135.8 | 1 | 0 | **0** |
| 1583.5 | 13 | 0 | **0** | 2776 | 1 | 0 | **0** |
| 1589.5 | 13 | 0 | **0** | 2337.9 | 1 | 0 | **0** |
| 2352.9 | 13 | 0 | **0** | 2477 | 1 | 0 | **0** |
| 1461.5 | 12 | 0 | **0** | 1972.8 | 1 | 0 | **0** |
| 2759.9 | 12 | 0 | **0** | 2467.9 | 1 | 0 | **0** |
| 2237.8 | 12 | 0 | **0** | 2424.9 | 1 | 0 | **0** |
| 2059.7 | 12 | 0 | **0** | 2247.7 | 1 | 0 | **0** |
| 1868.7 | 12 | 0 | **0** | 1300.6 | 1 | 0 | **0** |
| 1502.6 | 12 | 0 | **0** | 2571 | 1 | 0 | **0** |
| 1501.6 | 11 | 0 | **0** | 2678 | 1 | 0 | **0** |
| 1550.6 | 11 | 0 | **0** | 1548.6 | 1 | 0 | **0** |
| 1754.7 | 11 | 0 | **0** | 2426.9 | 1 | 0 | **0** |
| 1695.6 | 11 | 0 | **0** | 2791.9 | 1 | 0 | **0** |
| 3407.2 | 11 | 0 | **0** | 2557.9 | 1 | 0 | **0** |
| 1939.7 | 11 | 0 | **0** | 1998.8 | 1 | 0 | **0** |
| 1914.7 | 11 | 0 | **0** | 3156.2 | 1 | 0 | **0** |
| 1404.5 | 11 | 0 | **0** | 2555.8 | 1 | 0 | **0** |
| 1096.4 | 11 | 0 | **0** | 2322.8 | 1 | 0 | **0** |
| 2075.7 | 11 | 0 | **0** | 2336.8 | 1 | 0 | **0** |
| 1858.7 | 10 | 0 | **0** | 1694.5 | 1 | 0 | **0** |
| 1446.6 | 10 | 0 | **0** | 1826.7 | 1 | 0 | **0** |
| 2028.7 | 10 | 0 | **0** | 2704.1 | 1 | 0 | **0** |
| 2311.9 | 10 | 0 | **0** | 1462.5 | 1 | 0 | **0** |
| 2350.8 | 10 | 0 | **0** | 1909.7 | 1 | 0 | **0** |
| 1202.4 | 10 | 0 | **0** | 2642.9 | 1 | 0 | **0** |
| 1405.5 | 9 | 0 | **0** | 2070.7 | 1 | 0 | **0** |
| 1956.7 | 9 | 0 | **0** | 3433.2 | 1 | 0 | **0** |
| 1915.7 | 9 | 0 | **0** | 2408.9 | 1 | 0 | **0** |
| 2621.9 | 9 | 0 | **0** | 3519.2 | 1 | 0 | **0** |
| 1524.5 | 9 | 0 | **0** | 1809.7 | 1 | 0 | **0** |
| 877.3 | 9 | 0 | **0** | 1940.7 | 1 | 0 | **0** |
| 3084.1 | 9 | 0 | **0** | 4028.5 | 1 | 0 | **0** |
| 1503.6 | 9 | 0 | **0** | 2717.9 | 1 | 0 | **0** |
| 2093.7 | 8 | 0 | **0** | 2469 | 1 | 0 | **0** |
| 2922 | 8 | 0 | **0** | 2802 | 1 | 0 | **0** |
| 2239.8 | 8 | 0 | **0** | 2461.9 | 1 | 0 | **0** |
| 2612.9 | 8 | 0 | **0** | 3301.1 | 1 | 0 | **0** |
| 2620.9 | 8 | 0 | **0** | 2582.9 | 1 | 0 | **0** |
| 1551.6 | 7 | 0 | **0** | 2948.1 | 1 | 0 | **0** |

| | | | | | | | |
|---|---|---|---|---|---|---|---|
| 2879 | 7 | 0 | **0** | 1856.6 | 1 | 0 | **0** |
| 3406.2 | 7 | 0 | **0** | 3011.1 | 1 | 0 | **0** |
| 408.2 | 7 | 0 | **0** | 3008 | 1 | 0 | **0** |
| 1299.5 | 7 | 0 | **0** | 2774 | 1 | 0 | **0** |
| 2272.8 | 7 | 0 | **0** | 3299.3 | 1 | 0 | **0** |
| 1710.6 | 7 | 0 | **0** | 553.2 | 1 | 0 | **0** |
| 1793.7 | 7 | 0 | **0** | 2112.8 | 1 | 0 | **0** |
| 2273.8 | 7 | 0 | **0** | 2263.8 | 1 | 0 | **0** |
| 2458.8 | 6 | 0 | **0** | 1954.7 | 1 | 0 | **0** |
| 2457.8 | 6 | 0 | **0** | 2053.7 | 1 | 0 | **0** |
| 3081.1 | 6 | 0 | **0** | 2877.9 | 1 | 0 | **0** |
| 1097.4 | 6 | 0 | **0** | 2793 | 1 | 0 | **0** |
| 2775 | 6 | 0 | **0** | 2646 | 1 | 0 | **0** |
| 1810.7 | 6 | 0 | **0** | 3048.2 | 1 | 0 | **0** |
| 407.2 | 6 | 0 | **0** | 2587 | 1 | 0 | **0** |
| 2216.9 | 6 | 0 | **0** | 3116.1 | 1 | 0 | **0** |
| 2622.9 | 6 | 0 | **0** | 3009 | 1 | 0 | **0** |
| 1955.7 | 6 | 0 | **0** | 2483.9 | 1 | 0 | **0** |
| 714.3 | 6 | 0 | **0** | 2497.9 | 1 | 0 | **0** |
| 1852.6 | 6 | 0 | **0** | 2571.8 | 1 | 0 | **0** |
| 2207.7 | 6 | 0 | **0** | 716.3 | 1 | 0 | **0** |
| 2092.7 | 6 | 0 | **0** | 2274.9 | 1 | 0 | **0** |
| 1907.7 | 6 | 0 | **0** | 2880.1 | 1 | 0 | **0** |
| 2159.9 | 6 | 0 | **0** | 2314.9 | 1 | 0 | **0** |
| 1687.6 | 5 | 0 | **0** | 2574 | 1 | 0 | **0** |
| 1946.7 | 5 | 0 | **0** | 4029.5 | 1 | 0 | **0** |
| 2069.9 | 5 | 0 | **0** | 2278.7 | 1 | 0 | **0** |
| 2353.8 | 5 | 0 | **0** | 3230.2 | 1 | 0 | **0** |
| 2482.9 | 5 | 0 | **0** | 2175.8 | 1 | 0 | **0** |
| 2061.7 | 5 | 0 | **0** | 3050.1 | 1 | 0 | **0** |
| 2312.8 | 5 | 0 | **0** | 2391.9 | 1 | 0 | **0** |
| 1712.6 | 5 | 0 | **0** | 3405.2 | 1 | 0 | **0** |
| 2465.9 | 5 | 0 | **0** | 554.2 | 1 | 0 | **0** |
| 2619.9 | 5 | 0 | **0** | 2116.8 | 1 | 0 | **0** |
| 1663.6 | 5 | 0 | **0** | | | | |

**Supplementary Table 8**: FDR for each mass offset for O-glycan main search (110 glycans) from Yang *et al.*

| Modification Mass | Target PSMs | Decoy PSMs | **FDR %** | Modification Mass | Target PSMs | Decoy PSMs | **FDR %** |
|---|---|---|---|---|---|---|---|
| 365.1 | 44732 | 108 | **0.2** | 691.3 | 70 | 0 | **0** |
| 42 | 29805 | 8 | **0** | 1706.6 | 69 | 0 | **0** |
| 1 | 16519 | 12 | **0.1** | 2556.9 | 69 | 0 | **0** |
| 16 | 15841 | 1 | **0** | 1647.6 | 69 | 0 | **0** |
| 730.3 | 13232 | 35 | **0.3** | 919.3 | 68 | 0 | **0** |
| 57 | 11947 | 38 | **0.3** | 1243.5 | 68 | 0 | **0** |
| 80 | 10941 | 100 | **0.9** | 1362.5 | 67 | 2 | **3** |
| 366.1 | 9148 | 0 | **0** | 1046.4 | 67 | 0 | **0** |
| 0 | 6382 | 0 | **0** | 1151.4 | 66 | 2 | **3** |
| 203.1 | 5503 | 0 | **0** | 529.2 | 66 | 0 | **0** |
| 1095.4 | 5362 | 19 | **0.4** | 1907.7 | 66 | 0 | **0** |
| 367.1 | 4547 | 0 | **0** | 918.4 | 65 | 1 | **1.5** |
| 731.3 | 3349 | 2 | **0.1** | 1201.5 | 65 | 2 | **3.1** |
| 1096.4 | 2695 | 1 | **0** | 2030.7 | 65 | 0 | **0** |
| 568.2 | 2677 | 14 | **0.5** | 990.4 | 65 | 0 | **0** |
| 933.3 | 1510 | 4 | **0.3** | 1753.6 | 64 | 0 | **0** |
| 1461.5 | 1477 | 25 | **1.7** | 1542.5 | 63 | 0 | **0** |
| 732.3 | 1154 | 0 | **0** | 861.3 | 63 | 0 | **0** |
| 1460.5 | 1037 | 42 | **4.1** | 1987.7 | 63 | 0 | **0** |
| 771.3 | 883 | 6 | **0.7** | 1576.6 | 61 | 0 | **0** |
| 569.2 | 840 | 1 | **0.1** | 756.3 | 61 | 0 | **0** |
| 1097.4 | 829 | 0 | **0** | 1540.5 | 60 | 0 | **0** |
| 1462.5 | 755 | 4 | **0.5** | 1047.4 | 59 | 0 | **0** |
| 2135.8 | 750 | 0 | **0** | 1193.5 | 58 | 0 | **0** |
| 204 | 747 | 0 | **0** | 1590.6 | 57 | 0 | **0** |
| 1826.7 | 734 | 15 | **2** | 1079.4 | 57 | 0 | **0** |
| 406.2 | 733 | 1 | **0.1** | 2118.8 | 57 | 0 | **0** |
| 1827.7 | 710 | 4 | **0.6** | 1354.5 | 57 | 0 | **0** |
| 1136.4 | 686 | 7 | **1** | 1194.5 | 56 | 0 | **0** |
| 934.3 | 626 | 0 | **0** | 1534.5 | 55 | 0 | **0** |
| 2500.9 | 582 | 0 | **0** | 1704.6 | 55 | 1 | **1.8** |
| 1298.5 | 568 | 26 | **4.6** | 1397.6 | 55 | 0 | **0** |
| 1299.5 | 500 | 11 | **2.2** | 1282.5 | 54 | 1 | **1.9** |
| 2134.8 | 478 | 0 | **0** | 1013.4 | 54 | 0 | **0** |
| 408.1 | 463 | 0 | **0** | 1768.6 | 53 | 0 | **0** |
| 570.2 | 462 | 0 | **0** | 1355.5 | 53 | 0 | **0** |
| 1137.4 | 431 | 1 | **0.2** | 2029.7 | 53 | 0 | **0** |
| 205.1 | 405 | 1 | **0.2** | 1403.5 | 53 | 0 | **0** |
| 859.3 | 386 | 0 | **0** | 1309.5 | 52 | 0 | **0** |

| | | | | | | | |
|---|---|---|---|---|---|---|---|
| 511.2 | 360 | 0 | **0** | 2119.8 | 52 | 0 | **0** |
| 494.2 | 351 | 0 | **0** | 1386.5 | 52 | 0 | **0** |
| 772.3 | 327 | 1 | **0.3** | 1445.5 | 52 | 0 | **0** |
| 1989.7 | 312 | 3 | **1** | 2498.9 | 52 | 0 | **0** |
| 610.2 | 301 | 1 | **0.3** | 1040.4 | 51 | 2 | **3.9** |
| 1664.6 | 298 | 14 | **4.7** | 1202.5 | 51 | 2 | **3.9** |
| 2192.8 | 297 | 1 | **0.3** | 1356.5 | 51 | 0 | **0** |
| 1241.5 | 295 | 1 | **0.3** | 1727.6 | 51 | 0 | **0** |
| 1502.6 | 295 | 3 | **1** | 991.4 | 51 | 0 | **0** |
| 609.3 | 290 | 0 | **0** | 1866.7 | 50 | 0 | **0** |
| 876.3 | 286 | 0 | **0** | 1549.5 | 50 | 1 | **2** |
| 349.2 | 285 | 0 | **0** | 893.3 | 49 | 0 | **0** |
| 2558 | 280 | 0 | **0** | 1971.7 | 49 | 0 | **0** |
| 1501.6 | 277 | 9 | **3.2** | 1200.4 | 49 | 3 | **6.1** |
| 407.2 | 253 | 0 | **0** | 2175.8 | 49 | 0 | **0** |
| 1300.5 | 251 | 2 | **0.8** | 1023.3 | 49 | 0 | **0** |
| 1825.7 | 247 | 3 | **1.2** | 1226.5 | 49 | 0 | **0** |
| 1242.5 | 237 | 0 | **0** | 2703 | 49 | 0 | **0** |
| 1224.4 | 234 | 1 | **0.4** | 1218.5 | 48 | 0 | **0** |
| 1607.6 | 234 | 0 | **0** | 853.3 | 47 | 0 | **0** |
| 974.4 | 229 | 0 | **0** | 1687.6 | 46 | 0 | **0** |
| 935.4 | 223 | 0 | **0** | 1378.5 | 46 | 0 | **0** |
| 611.3 | 218 | 0 | **0** | 1589.6 | 45 | 0 | **0** |
| 1988.7 | 214 | 1 | **0.5** | 1005.3 | 45 | 0 | **0** |
| 2499.9 | 209 | 0 | **0** | 1906.7 | 45 | 0 | **0** |
| 1138.5 | 207 | 0 | **0** | 1524.5 | 44 | 0 | **0** |
| 513.2 | 206 | 0 | **0** | 2190.7 | 44 | 0 | **0** |
| 1663.6 | 206 | 4 | **1.9** | 852.3 | 43 | 0 | **0** |
| 716.3 | 201 | 0 | **0** | 1565.6 | 43 | 0 | **0** |
| 860.3 | 199 | 0 | **0** | 844.3 | 43 | 0 | **0** |
| 656.2 | 196 | 0 | **0** | 1323.5 | 43 | 3 | **7** |
| 1665.6 | 195 | 2 | **1** | 1591.6 | 42 | 0 | **0** |
| 1339.5 | 192 | 0 | **0** | 1283.5 | 41 | 0 | **0** |
| 1606.6 | 189 | 0 | **0** | 843.3 | 40 | 0 | **0** |
| 2354.9 | 189 | 0 | **0** | 1444.5 | 40 | 0 | **0** |
| 1340.5 | 189 | 0 | **0** | 1192.5 | 39 | 0 | **0** |
| 714.3 | 187 | 0 | **0** | 1080.4 | 39 | 0 | **0** |
| 1503.5 | 181 | 1 | **0.6** | 1259.5 | 39 | 0 | **0** |
| 2191.8 | 180 | 1 | **0.6** | 820.3 | 38 | 0 | **0** |
| 689.2 | 177 | 1 | **0.6** | 1379.5 | 37 | 0 | **0** |
| 512.2 | 172 | 1 | **0.6** | 2393.9 | 37 | 0 | **0** |
| 877.4 | 170 | 0 | **0** | 1575.6 | 37 | 0 | **0** |
| 351.1 | 168 | 0 | **0** | 2214.8 | 37 | 0 | **0** |

| | | | | | | | |
|---|---|---|---|---|---|---|---|
| 773.3 | 167 | 0 | **0** | 1014.3 | 37 | 0 | **0** |
| 2704 | 164 | 0 | **0** | 1648.6 | 36 | 0 | **0** |
| 495.1 | 161 | 0 | **0** | 1185.4 | 36 | 0 | **0** |
| 803.3 | 160 | 0 | **0** | 851.3 | 35 | 1 | **2.9** |
| 350.1 | 160 | 0 | **0** | 894.3 | 35 | 0 | **0** |
| 527.1 | 159 | 0 | **0** | 1446.6 | 35 | 0 | **0** |
| 715.3 | 155 | 0 | **0** | 1532.5 | 35 | 0 | **0** |
| 496.2 | 155 | 0 | **0** | 1007.4 | 34 | 0 | **0** |
| 641.3 | 149 | 0 | **0** | 1913.7 | 34 | 0 | **0** |
| 642.2 | 148 | 0 | **0** | 1220.5 | 34 | 0 | **0** |
| 1054.4 | 147 | 0 | **0** | 1364.5 | 34 | 0 | **0** |
| 786.3 | 140 | 0 | **0** | 1915.8 | 33 | 0 | **0** |
| 1153.4 | 139 | 0 | **0** | 1930.7 | 33 | 0 | **0** |
| 1623.6 | 138 | 0 | **0** | 1183.4 | 33 | 0 | **0** |
| 640.3 | 138 | 0 | **0** | 819.3 | 33 | 0 | **0** |
| 975.4 | 135 | 0 | **0** | 1387.5 | 32 | 0 | **0** |
| 1048.4 | 134 | 0 | **0** | 1308.5 | 32 | 0 | **0** |
| 755.3 | 134 | 0 | **0** | 1258.5 | 32 | 0 | **0** |
| 1341.5 | 131 | 0 | **0** | 1257.5 | 32 | 0 | **0** |
| 147.1 | 129 | 0 | **0** | 1914.6 | 32 | 1 | **3.1** |
| 802.3 | 129 | 0 | **0** | 1363.5 | 31 | 2 | **6.5** |
| 1769.6 | 127 | 0 | **0** | 1754.6 | 31 | 0 | **0** |
| 2133.8 | 126 | 0 | **0** | 1081.4 | 31 | 0 | **0** |
| 1152.4 | 126 | 0 | **0** | 1649.6 | 31 | 0 | **0** |
| 787.3 | 125 | 0 | **0** | 836.3 | 30 | 10 | **33.3** |
| 1972.7 | 124 | 0 | **0** | 1321.5 | 30 | 0 | **0** |
| 1021.4 | 124 | 0 | **0** | 1557.6 | 30 | 0 | **0** |
| 699.3 | 124 | 0 | **0** | 1525.6 | 30 | 0 | **0** |
| 1973.7 | 121 | 0 | **0** | 1216.4 | 30 | 0 | **0** |
| 1015.4 | 121 | 1 | **0.8** | 1380.5 | 30 | 0 | **0** |
| 813.4 | 120 | 0 | **0** | 2352.9 | 29 | 0 | **0** |
| 788.3 | 118 | 0 | **0** | 1688.6 | 29 | 0 | **0** |
| 997.3 | 118 | 1 | **0.8** | 1905.7 | 29 | 0 | **0** |
| 845.4 | 116 | 0 | **0** | 2556 | 29 | 0 | **0** |
| 2215.8 | 114 | 0 | **0** | 1548.5 | 29 | 0 | **0** |
| 2353.9 | 114 | 0 | **0** | 1396.6 | 28 | 0 | **0** |
| 1225.4 | 111 | 0 | **0** | 1161.4 | 28 | 0 | **0** |
| 804.3 | 111 | 0 | **0** | 2213.7 | 27 | 0 | **0** |
| 2176.8 | 111 | 0 | **0** | 1284.5 | 27 | 0 | **0** |
| 1868.7 | 107 | 0 | **0** | 1566.6 | 26 | 0 | **0** |
| 1017.3 | 107 | 0 | **0** | 1559.6 | 26 | 0 | **0** |
| 998.3 | 106 | 9 | **8.5** | 1160.4 | 25 | 0 | **0** |
| 1624.6 | 100 | 0 | **0** | 2093.7 | 25 | 0 | **0** |

| | | | |
|---|---|---|---|
| 658.2 | 99 | 0 | **0** |
| 1608.6 | 98 | 0 | **0** |
| 812.3 | 98 | 1 | **1** |
| 674.3 | 97 | 2 | **2.1** |
| 1752.7 | 96 | 0 | **0** |
| 1770.7 | 96 | 0 | **0** |
| 146 | 96 | 0 | **0** |
| 673.2 | 93 | 0 | **0** |
| 690.3 | 93 | 0 | **0** |
| 698.3 | 92 | 0 | **0** |
| 528.2 | 91 | 0 | **0** |
| 878.3 | 91 | 0 | **0** |
| 697.3 | 91 | 0 | **0** |
| 757.3 | 90 | 0 | **0** |
| 1016.4 | 90 | 1 | **1.1** |
| 1622.6 | 87 | 0 | **0** |
| 675.3 | 87 | 0 | **0** |
| 1832.6 | 87 | 0 | **0** |
| 1831.6 | 83 | 3 | **3.6** |
| 1039.3 | 83 | 7 | **8.4** |
| 892.3 | 81 | 0 | **0** |
| 1038.4 | 81 | 0 | **0** |
| 657.2 | 80 | 0 | **0** |
| 814.4 | 79 | 0 | **0** |
| 989.4 | 78 | 0 | **0** |
| 917.4 | 78 | 0 | **0** |
| 976.4 | 77 | 0 | **0** |
| 1867.7 | 77 | 0 | **0** |
| 148.1 | 76 | 0 | **0** |
| 1022.4 | 76 | 0 | **0** |
| 818.3 | 75 | 0 | **0** |
| 1006.4 | 74 | 0 | **0** |
| 1055.4 | 73 | 0 | **0** |
| 1705.6 | 73 | 0 | **0** |
| 999.3 | 72 | 3 | **4.2** |
| 1686.6 | 72 | 0 | **0** |
| 2092.8 | 72 | 0 | **0** |
| 835.3 | 71 | 1 | **1.4** |
| 1541.5 | 71 | 0 | **0** |
| 1008.4 | 71 | 0 | **0** |
| 700.3 | 71 | 0 | **0** |

| | | | |
|---|---|---|---|
| 2395.9 | 24 | 0 | **0** |
| 1532.6 | 23 | 0 | **0** |
| 1574.6 | 23 | 0 | **0** |
| 1405.5 | 22 | 0 | **0** |
| 1056.4 | 22 | 0 | **0** |
| 1558.6 | 21 | 0 | **0** |
| 2174.8 | 21 | 0 | **0** |
| 1217.4 | 21 | 0 | **0** |
| 1219.5 | 21 | 0 | **0** |
| 2117.8 | 20 | 0 | **0** |
| 1728.6 | 20 | 0 | **0** |
| 837.3 | 20 | 0 | **0** |
| 1916.7 | 20 | 0 | **0** |
| 1931.7 | 20 | 0 | **0** |
| 1307.5 | 19 | 1 | **5.3** |
| 2018.7 | 19 | 0 | **0** |
| 1395.5 | 19 | 0 | **0** |
| 2702 | 19 | 0 | **0** |
| 1573.6 | 18 | 0 | **0** |
| 1533.6 | 18 | 0 | **0** |
| 1184.4 | 18 | 0 | **0** |
| 1322.5 | 17 | 0 | **0** |
| 1388.5 | 17 | 0 | **0** |
| 1526.6 | 17 | 0 | **0** |
| 1567.6 | 17 | 0 | **0** |
| 1833.7 | 16 | 0 | **0** |
| 1550.6 | 16 | 0 | **0** |
| 1729.6 | 15 | 0 | **0** |
| 1548.6 | 15 | 0 | **0** |
| 2094.7 | 15 | 0 | **0** |
| 1932.7 | 13 | 0 | **0** |
| 1243.4 | 12 | 0 | **0** |
| 2028.7 | 12 | 0 | **0** |
| 2394.8 | 11 | 0 | **0** |
| 2020.8 | 11 | 0 | **0** |
| 1159.5 | 8 | 0 | **0** |
| 1542.6 | 7 | 0 | **0** |
| 1395.6 | 5 | 0 | **0** |
| 2019.7 | 4 | 0 | **0** |
| 1589.5 | 3 | 0 | **0** |
